# Inter-subject correlations and their behavioral associations vary across movies: Implications for generalizability

**DOI:** 10.1101/2024.12.03.626542

**Authors:** Simon Leipold, Rajat Ravi Rao, Jan-Mathijs Schoffelen, Sara Bögels, Ivan Toni

## Abstract

Movie-watching fMRI has become increasingly popular in neuroscience. Movie-fMRI data are commonly analyzed using inter-subject correlation (ISC), which quantifies the similarity of neural time series across individuals. Differences in ISC during movie viewing have been associated with psychological traits and clinical diagnoses. However, most studies investigating group differences in ISC or ISC–behavior associations have drawn conclusions from a single movie. Because ISC is inherently stimulus-driven, effects observed for one movie may not generalize to another. Yet, the extent to which ISC patterns and ISC–behavior associations depend on the specific movie being viewed has received limited systematic attention.

Here, we analyzed three independent datasets comprising 318 subjects and 36 movies in total to quantify between-movie variability in ISC and assess its consequences for ISC–behavior associations. Across datasets, ISC varied between movies throughout the cortex. This variability was spatially heterogeneous: regions with stronger ISC showed greater between-movie variability. Movie-specific inter-subject representational similarity analysis revealed distinct spatial distributions of ISC–behavior associations, with limited overlap between movies. This pattern was observed for two distinct behavioral constructs. These findings suggest that ISC–behavior associations can be strongly movie-specific.

## 1. Introduction

Over the past two decades, movie-watching paradigms have become increasingly popular in neuroscience (Eickhoff et al., 2020; Hasson et al., 2010; Jääskeläinen et al., 2021; Redcay & Moraczewski, 2020; Sonkusare et al., 2019; Vanderwal et al., 2019). In these paradigms, subjects view movies while undergoing neuroimaging, most commonly functional magnetic resonance imaging (fMRI). A common approach for analyzing fMRI data acquired during movie watching is inter-subject correlation (ISC) analysis (Hasson et al., 2004). In ISC analysis, the fMRI time series from a given brain region, such as a voxel or atlas parcel, in one subject is used as a model for the fMRI time series from the same region in another subject (Nastase et al., 2019). The resulting correlation coefficient quantifies the similarity, or synchronization, of neural activity between subjects (Nummenmaa et al., 2018). ISC can be computed pairwise across all possible subject pairs or by correlating each subject’s time series with the group-average time series. The resulting correlations are then typically summarized, for example by averaging across subject pairs, to estimate the overall level of synchronization within each brain region. These group-level estimates are themselves commonly referred to as ISC.

Prior work has demonstrated that movies evoke strong ISC across the cortex, with particularly strong synchronization in visual and auditory brain regions (Hasson et al., 2004; Nummenmaa et al., 2018). An advantage of ISC over general linear model analysis is that it is data-driven and does not require explicit regressors for specific event types, such as the appearance of faces or objects during the movie. Crucially, ISC is *stimulus-driven*: activation changes that are not consistently related to processing of the movies across subjects remain largely uncorrelated and do not contribute to ISC estimates (Nastase et al., 2019; Nummenmaa et al., 2018). This stimulus sensitivity is a key strength of ISC, as it allows researchers to identify brain responses that are reliably driven by shared stimulus processing. However, the extent to which different movies evoke similar or distinct spatial patterns of ISC has rarely been examined systematically.

In recent years, studies using movie-fMRI paradigms have increasingly examined group differences in ISC. In a typical study, ISC is compared between groups or subgroups of subjects defined by characteristics such as psychological traits or clinical diagnoses. For example, friends show higher ISC than individuals who are more distant from one another in a social network (Parkinson et al., 2018). Similarly, sisters show higher ISC than friends (Bacha-Trams et al., 2024), and married couples show higher ISC than randomly paired individuals (L. Li et al., 2022). In the clinical domain, studies have reported altered ISC during movie watching in individuals with autism (Hasson et al., 2009; Mangnus et al., 2024; Salmi et al., 2013), psychosis (Mäntylä et al., 2018), depression (Guo et al., 2015), and social anxiety (Camacho et al., 2023; Koch et al., 2026), among other clinical populations. This line of research has recently been extended through inter-subject representational similarity analysis (IS-RSA), which enables a dimensional assessment of the association between subjects’ neural synchronization patterns and behavioral characteristics (Finn et al., 2020; Nummenmaa et al., 2012). For example, subjects who are more similar in personality traits (Matz et al., 2022) or political orientation (de Bruin et al., 2023; Katabi et al., 2023; Leong et al., 2020; van Baar et al., 2021) show higher ISC during movie watching than subjects who are less similar in these characteristics.

Most studies investigating group differences in ISC or associations between ISC and behavior have drawn conclusions from a single film or movie clip. However, the few studies using multiple movies suggest that different movies can evoke distinct spatial patterns of neural synchronization across the brain (Camacho et al., 2023; Katabi et al., 2023; L. Li et al., 2022; Sievers et al., 2024; van Baar et al., 2021). This is plausible because movies are spatially and temporally complex, highly multidimensional stimulus sequences that vary across numerous perceptual, narrative, affective, and stylistic features (Bartels & Zeki, 2004; Hasson et al., 2004). Indeed, the sensitivity of ISC to such features was already highlighted in the earliest ISC studies (Hasson et al., 2004; Hasson, Landesman, et al., 2008; Hasson et al., 2010). Because ISC is inherently stimulus-driven (Nastase et al., 2019; Nummenmaa et al., 2018), group differences or ISC–behavior associations observed for one movie may not necessarily generalize to another. However, the extent and interpretational consequences of this movie-specificity have received limited systematic attention.

The extent to which ISC findings generalize from one movie to another is central for defining the scope of conclusions drawn from studies using one or a few movies (Sonkusare et al., 2019; Vanderwal et al., 2019). We focus on generalizability across stimulus materials, defined here as the extent to which findings obtained with one movie would be expected to recur for other movies that could have been used to address the same research question (Yarkoni, 2022). This does not imply that findings from one movie should generalize to all possible movies. Rather, the question is how strongly ISC findings depend on the movie selected, and whether they remain stable across other movies. Generalizability across movies is therefore a graded rather than all-or-none property. At one extreme, an ISC-based finding may generalize across many different movies; at the other, it may be specific to the particular movie used in a study.

Here, we leverage three independent datasets, comprising one discovery dataset and two replication datasets, with 36 movies and 318 subjects in total, to investigate how ISC and ISC–behavior associations vary across movies (**Figure 1**). In the discovery dataset, 112 adults each watched eight animated short films (Eijk et al., 2022) and completed a task assessing conceptual judgments of novel objects, which we related to ISC using IS-RSA. In the first replication dataset, 178 subjects watched 14 short live-action movie clips (Van Essen et al., 2013); in the second replication dataset, 28 subjects watched 14 live-action and animated short films (Morgenroth et al., 2025). For both replication datasets, we related similarity between subjects in Big Five personality traits (Goldberg, 1993) to ISC. Across the three datasets, ISC varied between movies across the cortex, with the extent of between-movie variability differing across brain regions: regions with stronger ISC showed greater variability. Movie-specific IS-RSA analyses revealed strongly varying ISC–behavior associations across movies, a pattern observed for both behavioral constructs.

**Figure 1.**
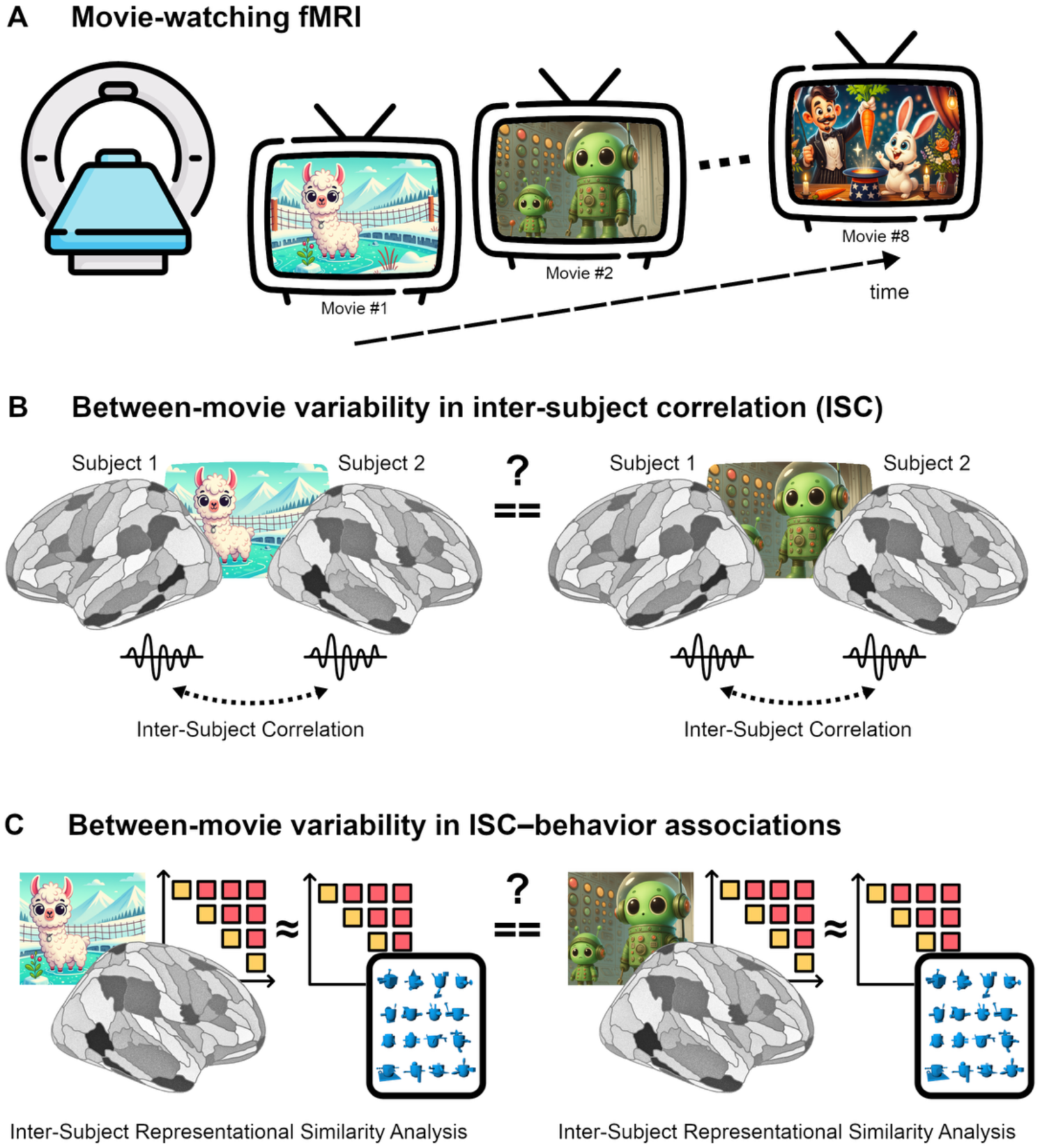
Overview. **A)** In the CABB dataset, subjects (*N* = 112) viewed eight animated movies during fMRI scanning. The movies were selected to include diverse object categories, such as humans, plants, tools, toys, and food. **B)** We quantified the variability in inter-subject correlation (ISC) between movies across the brain. For each subject and movie, we extracted a spatially averaged fMRI time series from 210 cortical parcels. ISC was computed for each parcel by correlating the time series between all subject pairs. **C)** To assess the consequences of between-movie variability for ISC–behavior associations, we used inter-subject representational similarity (IS-RSA) to relate neural synchronization during movie watching to the similarity of subjects’ conceptual representations of novel objects (“Fribbles”, in blue), with analyses conducted separately for each movie.

## 2. Methods

### 2.1. Discovery dataset: Communicative Alignment of Brain and Behaviour (CABB)

#### 2.1.1. Subjects

We analyzed data from *N* = 112 subjects from the Communicative Alignment of Brain and Behaviour (CABB) dataset (Eijk et al., 2022). These subjects were the same as those included in the fMRI analysis in the dataset description paper by Eijk et al. (2022) and did not include subjects who were excluded due to technical malfunctions, incomplete data, or poor data quality. Subjects had a mean age of 22.72 years (standard deviation = 3.21 years); 36 were male and 76 were female. All subjects reported no history of neurological or language-related disorders, no brain surgery, no hearing impairments, no metal implants other than dental implants, and normal or corrected-to-normal vision. Written informed consent was obtained from all subjects for being included in the study. For our main analysis, a repeated-measures ANOVA on ISC values of 56 subject pairs, we had 80% statistical power to detect differences in ISC between movies with an effect size of Cohen’s *f* > 0.51. Statistical power was calculated using the *WebPower* package (version 0.9.2) (Zhang & Yuan, 2018) in *R* (version 4.1.0; RRID:SCR_001905).

#### 2.1.2. fMRI movie tasks

Subjects viewed eight animated movies (see **Supplementary Methods** for details), which were presented using Presentation software (version 20.2; RRID:SCR_002521). Each movie was a complete animated short film with a self-contained narrative, rather than an excerpt from a longer film. The movies were selected to include categories of objects frequently mentioned in pilot tests of novel objects (“Fribbles”, **Figure 1C**) used as referents by the subjects during face-to-face dialogues, such as humans, plants, tools, toys, and food. These animated movies are not geared exclusively toward children, but are accessible and engaging for a general audience, including adults. The movies were presented on a portion of the screen slightly below the center, on a black background with a height of 360 pixels and a width of 640 pixels, with a visual angle of approximately 9–11° vertically and 16–20° horizontally. The movies were played at 30 frames per second. Each movie was played in its entirety, except for the start and end (i.e., titles and credits), which were cut off to ensure no text was shown to subjects. The average duration of the movies was 4.1 minutes (range = 2.2–6.1 minutes), totaling 35 minutes, including 12-second breaks between subsequent movies. The movies were preceded by a filler video clip that lasted a few seconds. Subjects were instructed to simply attend to the movies and to lie still. The movies did not contain spoken language, but the original soundtrack was provided to subjects via earphones. Before scanning began, the sound levels were adjusted to each subject’s individual preferences.

#### 2.1.3. Imaging data acquisition

Magnetic resonance images were acquired using two 3T MAGNETOM magnetic resonance imaging scanners: Prisma and PrismaFit (Siemens AG, Healthcare Sector, Erlangen, Germany). Blood oxygenation level dependent (BOLD) functional images were acquired using a multi-band 2D-echo planar imaging (EPI) sequence (Uğurbil et al., 2013) with repetition time (TR) = 1000 ms, echo time (TE) = 34 ms, flip angle = 60°, 2 mm isotropic resolution, field of view (FOV) = 208x208 mm. A multi-band acceleration factor of 6 was used in the slice direction, and no parallel imaging was applied in-plane. Phase encoding was applied in the anterior-posterior (AP) direction with a partial Fourier coverage of 7/8. About 2074 volumes were acquired during movie watching. Anatomical images were acquired with a T1-weighted 3D MPRAGE sequence, integrated parallel acquisition technique (iPAT) acceleration factor = 2, with TR = 2200 ms, TE = 2 ms, inversion time (TI) = 1100 ms, flip angle = 11°, 0.8 mm isotropic resolution, FOV = 256x240x166 mm; and with a T2-weighted turbo spin echo (TSE) sequence with variable flip angle, TR = 3200 ms, TE = 569 ms, echo spacing = 4.12 ms, turbo factor = 314, 0.8 mm isotropic resolution, and FOV = 256x240x166 mm. Raw imaging data were converted to Brain Imaging Data Structure (BIDS) format (Gorgolewski et al., 2016) using *BIDScoin* (version 1.5; RRID:SCR_022839) (Zwiers et al., 2022).

#### 2.1.4. Imaging data preprocessing

Each subject’s raw imaging data underwent a quality control assessment for excessive motion, susceptibility artifacts, or other sources of noise using *MRIQC* (version 23.1.0; RRID:SCR_022942). Imaging data preprocessing was implemented using *fMRIPrep* (version 20.2.7; RRID:SCR_016216) (Esteban et al., 2019), which is based on *Nipype* (version 1.7.0; RRID:SCR_002502) (Gorgolewski et al., 2011). The main preprocessing steps included segmentation, surface reconstruction, and spatial normalization of the T1-weighted anatomical image; and slice timing correction, susceptibility distortion correction, head-motion correction, and spatial normalization of the fMRI data. Details of the preprocessing pipeline are included in the **Supplementary Methods**.

#### 2.1.5. fMRI data denoising

Each subject’s preprocessed fMRI data underwent further denoising, implemented using *Nilearn* (version 0.9.2; RRID:SCR_001362) in *Python* (version 3.7.13; RRID:SCR_008394). The voxel-wise fMRI time series were detrended and low-pass filtered at 0.08 Hz. Next, the following confounds estimated by fMRIprep during preprocessing were regressed out from the voxel-wise time series: six motion parameters (three translations, three rotations) and their first temporal derivatives, as well as cerebrospinal fluid and white matter time series and their first temporal derivatives. In addition, scans that exceeded a threshold of 0.5 mm framewise displacement (FD) or 1.5 standardized DVARS were annotated as motion outliers and included as nuisance regressors. Subsequently, the time series were standardized to have a mean of zero and unit variance. After denoising, we segmented the fMRI time series and extracted a separate time series for each movie based on the onset and offset times recorded in the Presentation log files. The denoised fMRI data for each movie was then spatially smoothed with a 6 mm full-width-at-half-maximum (FWHM) three-dimensional Gaussian kernel.

#### 2.1.6. Inter-subject correlation (ISC) analysis

To calculate parcel-wise ISC values, we first extracted a spatially averaged time series for each subject and movie from the cortical parcels of the Brainnetome atlas (Fan et al., 2016) using *NLtools* (version 0.4.7) (Chang et al., 2022) in *Python*. We restricted the analyses to the 210 cortical parcels of the Brainnetome atlas, excluding the 36 subcortical parcels, because the IS-RSA focused on conceptual representations of novel objects, which we expected to be primarily reflected in cortical synchronization patterns. For each movie and cortical parcel, we then calculated the ISC values for all possible subject pairs by computing the Pearson correlation coefficient between the parcel-wise fMRI time series of subject [i] and subject [j] of each pair [i,j] using *scikit-learn* (version 1.0.2; RRID:SCR_002577) in *Python*. To obtain a mean ISC value for each parcel, we averaged the ISC values across all subject pairs. Visualization of the ISC values in the brain was implemented using *Nilearn*.

#### 2.1.7. Statistical analyses of ISC values

Prior to statistical analysis, all ISC values were transformed using Fisher’s r-to-z transformation to normalize the distribution of correlation coefficients. To quantify between-movie variability in ISC across the whole cortex, we averaged the ISC values across all cortical parcels to obtain a single ISC value per subject pair and movie. Then, we entered the whole-brain ISC values into a repeated-measures ANOVA with Movie as the within-subject factor. We used data from 56 subject pairs, ensuring that each subject contributed only to a single ISC value per movie and that the assumptions of the repeated-measures ANOVA were not violated. Next, we repeated the ANOVA for each parcel separately to obtain a spatial distribution of between-movie variability across the cortex. The ANOVAs were performed using the *ez* (version 4.4-0; RRID:SCR_020990) and *rstatix* (version 0.7.2; RRID:SCR_021240) packages in *R*.

To investigate whether brain regions with high levels of ISC also show high variability in ISC, we correlated the parcel-wise level of ISC with the parcel-wise between-movie variability in ISC. Specifically, we extracted the *F* values from the repeated-measures ANOVAs for each parcel as a measure of between-movie variability. We then calculated the Pearson correlation coefficient between the *F* values and the ISC values for each parcel in *R*.

#### 2.1.8. Inter-subject representational similarity analysis (IS-RSA)

The CABB dataset encompasses behavioral data on subjects’ conceptual representations of novel objects (“Fribbles”, **Figure 1C**), which were estimated in a *Features task* completed before the movie-fMRI tasks (Binder et al., 2016; Eijk et al., 2022). In this task, subjects used a linear visual analog scale to report how well each novel object matched 29 different features (e.g., pointy, symmetrical, human, related to movement; full list in **Supplementary Table 1**; see **Supplementary Methods** for details on the Features task). Subjects showed substantial individual differences in these feature ratings, indicating variability in how the novel objects were conceptually represented across individuals (**Supplementary Figure 1**). Prior work has shown that subjects with similar interpretations of a narrative show higher neural synchronization than subjects with different interpretations (Yeshurun et al., 2017). Furthermore, subjects who share their interpretations of an ambiguous animation of geometric shapes show higher neural synchronization than those with dissimilar interpretations (Nguyen et al., 2019). Inspired by these previous studies, we hypothesized that subjects with similar conceptual representations of novel objects would show higher ISC during movie watching. As these particular movies were selected to contain categories of objects that were mentioned often in pilot tests to be associated with the novel objects (Eijk et al., 2022), we anticipated a “spill-over” effect from representations of novel objects to naturalistic viewing of conventional objects.

We conducted a regression-based IS-RSA (Finn et al., 2020; Nummenmaa et al., 2012) using linear mixed-effects models (G. Chen et al., 2017; van Baar et al., 2021) to link subject similarity in conceptual representations of the novel objects, estimated in the *Features task*, with neural synchronization during movie watching, separately for each movie. The linear mixed-effects models were estimated using the *lme4* package (version 1.1-33; RRID:SCR_015654) in *R* (version 4.1.0; RRID:SCR_001905). We estimated a full model to simultaneously assess the main effects of subjects’ similarities in conceptual representations on their ISC during movie watching, while controlling for subject similarity in lexical representations of the novel objects, estimated in a *Naming task*, and also controlling for subject similarities in sex and age. Details on the novel object stimuli, Features and Naming tasks, and the IS-RSA are given in the **Supplementary Methods**.

To ensure that our results were not driven by statistical thresholding, we additionally calculated Pearson correlations between all pairwise combinations of movie-specific, unthresholded IS-RSA *t*-maps and computed the average correlation separately for each dataset.

#### 2.1.9. Control analyses

We ran multiple additional analyses in the CABB dataset to test the robustness of our findings; these analyses are detailed in the **Supplementary Results**. Specifically, we (1) repeated all ISC and IS-RSA analyses using the Schaefer 300-parcel atlas (Schaefer et al., 2018) instead of the Brainnetome atlas, to ensure that the results were not dependent on the specific anatomical parcellation scheme used in the main analyses; (2) repeated all ISC and IS-RSA analyses using truncated time series to control for differences in movie length; (3) implemented a resampling approach to assess the stability of between-movie ISC variability across different non-overlapping subject-pair configurations, thereby testing whether the observed effects were robust to the particular pairing of subjects; (4) classified movies based on multivariate ISC patterns, to determine whether movie-specific ISC profiles were sufficiently distinct to identify individual movies from their spatial synchronization patterns; (5) controlled for differences in head motion across movies, to rule out the possibility that movie-specific differences in ISC were driven by systematic variation in subject motion; (6) controlled for parcel-wise temporal signal-to-noise ratio, to assess whether the association between overall ISC level and between-movie ISC variability could be explained by regional differences in data quality; (7) incorporated a random effect for movie in the IS-RSA model to evaluate whether any effects generalized across the movie set; and (8) repeated the IS-RSA analyses without covariates to confirm that the main findings were not dependent on the inclusion of control variables in the regression model.

### 2.2. Replication dataset 1: Human Connectome Project (HCP)

#### 2.2.1. Subjects

We analyzed data from the Human Connectome Project (HCP) dataset, S1200 release (Van Essen et al., 2013). After excluding six subjects with incomplete movie-watching data, the final sample included 178 subjects (mean age = 29.40 years, standard deviation = 3.31 years; 108 female, 70 male). The HCP study was approved by the Washington University Institutional Review Board, and all subjects provided informed consent.

#### 2.2.2. fMRI movie tasks

The HCP movie-watching protocol comprised two scanning sessions, each consisting of two runs of approximately 15 minutes. Each run included three to four short movie clips, separated by 20-second rest blocks, followed by a repeated validation clip. We excluded the validation clips because they consisted of concatenated brief clips lasting 1–3 seconds, resulting in 14 movie clips for analysis. Each analyzed clip was presented once. The analyzed clips had an average duration of 3.43 minutes, ranging from 1.06 to 4.30 minutes. Clips in the first and third runs were taken from independently produced Creative Commons films, whereas clips in the second and fourth runs were taken from Hollywood movies. Details on the movie clips are provided in the **Supplementary Methods**. All movie clips are available from the HCP website (https://db.humanconnectome.org/).

#### 2.2.3. Imaging data acquisition and preprocessing

Functional images were acquired on a 7T Siemens MAGNETOM scanner using a gradient-echo EPI sequence (Uğurbil et al., 2013) with TR = 1000 ms, TE = 22.2 ms, flip angle = 45°, 1.6 mm isotropic resolution, FOV = 208 x 208 mm, and a multiband acceleration factor of 5. We used the minimally preprocessed HCP data obtained from the HCP website. These data were processed with the standard HCP pipeline, including motion correction, spatial normalization, high-pass temporal filtering, regression of 24 motion parameters, and denoising (Glasser et al., 2013). We then segmented the preprocessed fMRI time series and extracted separate time series for each movie clip.

#### 2.2.4. ISC and IS-RSA analyses

As in the CABB dataset, we computed parcel-wise ISC values by first extracting a spatially averaged time series for each subject and movie clip from the cortical parcels of the Brainnetome atlas, using *NLtools* (version 0.5.1) in *Python* (version 3.11.15). For each movie clip and cortical parcel, we then computed ISC values for all possible subject pairs as the Pearson correlation between the corresponding parcel-wise fMRI time series. All ISC values were Fisher transformed to normalize the distribution of correlation coefficients. Statistical analyses comparing ISC values across movie clips were identical to those used for the CABB dataset. For the repeated-measures ANOVA, we used data from 89 randomly sampled, non-overlapping subject pairs. We also correlated parcel-wise ISC levels with parcel-wise between-movie ISC variability, following the same procedure as in the CABB dataset.

The HCP dataset includes behavioral data on subjects’ Big Five personality traits, based on the well-established Five-Factor Model (Goldberg, 1993; Heine & Buchtel, 2009). Personality traits were assessed using the 60-item version of the Five-Factor Inventory, which has shown excellent reliability and validity (McCrae & Costa, 2004). To obtain the subject-by-subject matrix for IS-RSA, we computed the correlation distance between each pair of subjects based on their scores across the five personality traits: openness to experience (NEOFAC_O), agreeableness (NEOFAC_A), conscientiousness (NEOFAC_C), extraversion (NEOFAC_E), and neuroticism (NEOFAC_N). As in the CABB dataset, we conducted regression-based IS-RSA using linear mixed-effects models to link subject similarity in personality traits with neural synchronization during movie watching, separately for each movie clip. The linear mixed-effects models were estimated using the *lme4* package (version 1.1-35) in *R* (version 4.3.3).

### 2.3. Replication dataset 2: Emotion Research using Films and fMRI (Emo-FilM)

#### 2.3.1. Subjects

We analyzed data from the Emotion Research using Films and fMRI (Emo-FilM) dataset (Morgenroth et al., 2025). The Emo-FilM fMRI dataset included 30 healthy subjects. We excluded two subjects because of missing data, yielding a final sample of 28 subjects (mean age = 25.32 years, standard deviation = 3.44 years; 17 female, 11 male). All subjects were right-handed, had normal or corrected-to-normal vision, full color vision, high English comprehension, no history of neurological or psychiatric conditions, and reported no use of neuropharmacological or recreational drugs. The study was approved by the Geneva Cantonal Commission for Ethics in Research (protocol No. 2018-02006), complied with the Code of Human Research Ethics (2014), and all subjects provided written informed consent.

#### 2.3.2. fMRI movie tasks

The Emo-FilM protocol comprised four fMRI sessions of approximately two hours each. Across these sessions, subjects watched 14 short films in pseudo-random order, with two to five films presented per session. The stimuli consisted of full live-action and animated short films with complete narratives and spoken language. The films were presented without beginning or end credits and had an average duration of 11.44 minutes, ranging from 6.70 to 17.14 minutes. Subjects were instructed to watch the films as they would in everyday life. Details on the films are provided in the **Supplementary Methods**.

#### 2.3.3. Imaging data acquisition and preprocessing

Functional images were acquired on a 3T Siemens MAGNETOM TIM Trio scanner using a 32-channel head coil and a gradient-echo EPI sequence: TR = 1300 ms, TE = 30 ms, flip angle = 64°, multiband acceleration factor = 3, 2.5 mm isotropic resolution, and FOV = 210 × 210 mm. We used preprocessed fMRI data downloaded from *OpenNeuro* (RRID:SCR_005031) via *DataLad* (RRID:SCR_003931) (Halchenko et al., 2021). These data were preprocessed separately for each movie. Preprocessing included motion correction, brain extraction, spatial normalization, spatial smoothing with a 6 mm FWHM Gaussian kernel, grand-mean intensity normalization, and high-pass temporal filtering. White matter and cerebrospinal fluid time courses were regressed out together with six motion parameters.

#### 2.3.4. ISC and IS-RSA analyses

We computed parcel-wise ISC values by first extracting a spatially averaged time series for each subject and movie from the cortical parcels of the Brainnetome atlas. For each movie and cortical parcel, we then computed ISC values for all possible subject pairs as the Pearson correlation between the corresponding parcel-wise fMRI time series. All ISC values were transformed using Fisher’s r-to-z transformation. Statistical analyses comparing ISC values across movies were identical to those used for the CABB dataset. For the repeated-measures ANOVA, we used data from 14 randomly sampled, non-overlapping subject pairs. We also correlated parcel-wise ISC levels with parcel-wise between-movie ISC variability, following the same procedure as in the CABB dataset.

The Emo-FilM dataset includes behavioral data on subjects’ Big Five personality traits, based on the well-established Five-Factor Model (Goldberg, 1993; Heine & Buchtel, 2009). Personality traits were assessed using the Big Five Inventory (John et al., 1991). To construct the subject-by-subject matrix for IS-RSA, we computed the correlation distance between each pair of subjects based on their scores across the five personality traits: openness to experience (BIG5_ope), agreeableness (BIG5_agr), conscientiousness (BIG5_con), extraversion (BIG5_ext), and neuroticism (BIG5_neu).

As in the CABB dataset, we conducted regression-based IS-RSA using linear mixed-effects models to link subject similarity in personality traits with neural synchronization during movie watching, separately for each movie. The linear mixed-effects models were estimated using the *lme4* package (version 1.1-35) in *R* (version 4.3.3).

## 3. Results

### 3.1. ISC patterns vary between movies

Our first aim was to quantify variability in ISC evoked by different movies across the cortex. We analyzed fMRI data from 112 subjects from the CABB dataset (Eijk et al., 2022), acquired while subjects watched eight animated short films.

As shown in **Figure 2**, the overall spatial distribution of ISC evoked by each of the eight movies resembled the patterns reported in previous movie-fMRI studies (Hasson et al., 2004, 2010; Vanderwal et al., 2019). The strongest ISC was observed in the extended visual system, including face-sensitive regions in the fusiform gyrus, and in the auditory system, including Heschl’s gyrus and the planum temporale. Strong ISC was also observed in higher-order temporal, parietal, and frontal cortices.

**Figure 2.**
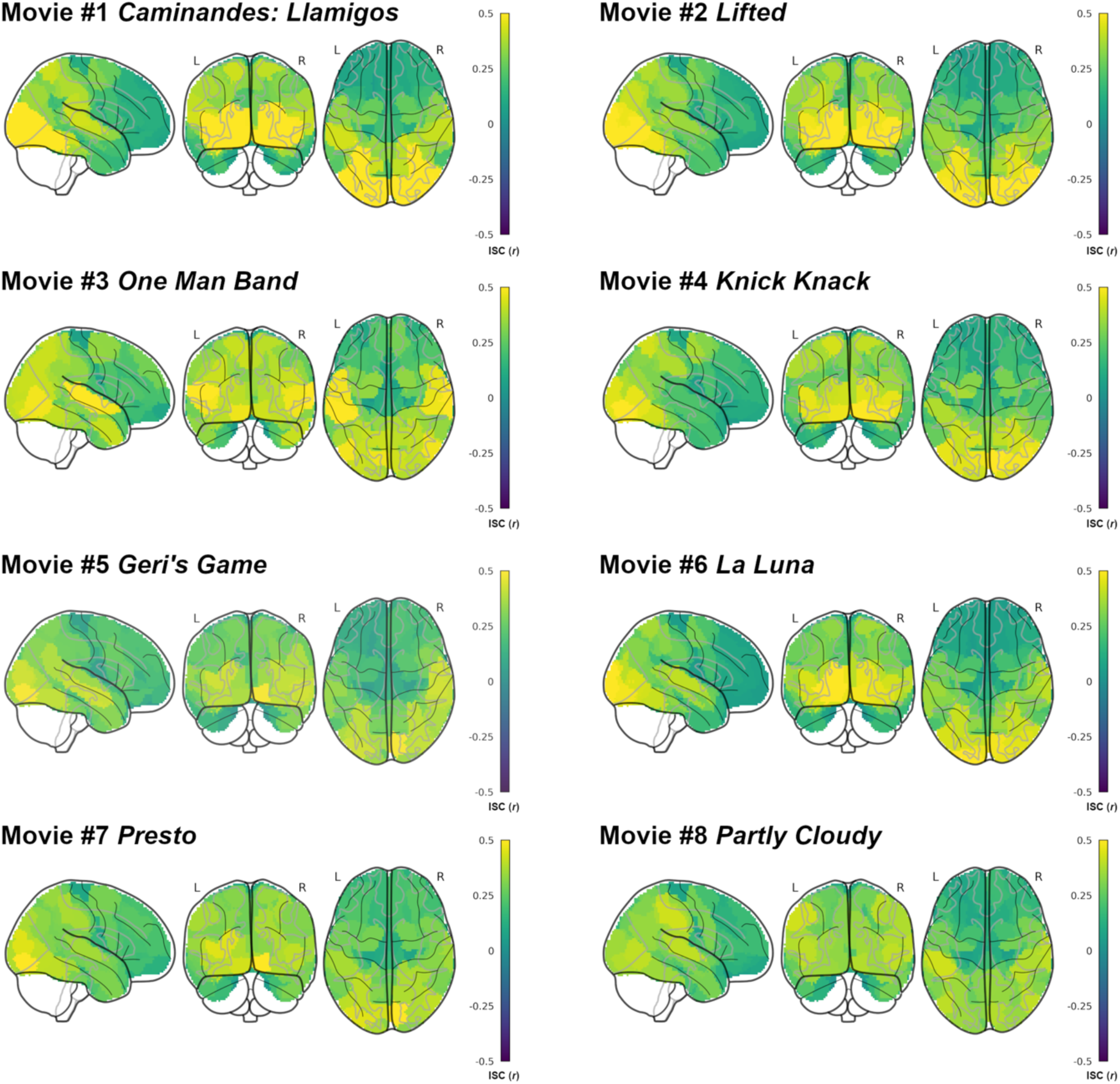
Inter-subject correlations for individual movies in the CABB dataset. Spatial distribution and magnitude of inter-subject correlations (ISC) across the cortex, shown separately for each of the eight movies.

We next quantified between-movie variability in ISC statistically. First, we tested whether whole-brain ISC varied between movies. To this end, we averaged ISC values across all cortical parcels to obtain one whole-brain ISC value per subject pair and movie. We then submitted the whole-brain ISC values to a repeated-measures analysis of variance (ANOVA) with Movie as the within-subject factor. To meet the assumptions of the ANOVA, we used a subset of non-overlapping subject pairs, ensuring that each subject contributed to only one pair. The ANOVA revealed a significant main effect of Movie (*F*(7,385) = 4.93, *p* < 0.001, generalized eta-squared [η^2^_G_] = 0.051), indicating considerable variability in whole-brain ISC across movies. **Figure 3A** shows the mean and standard error of whole-brain ISC for each movie. The differences in ISC were not driven by a single movie differing from all others; rather, whole-brain ISC varied across the full set of movies.

**Figure 3.**
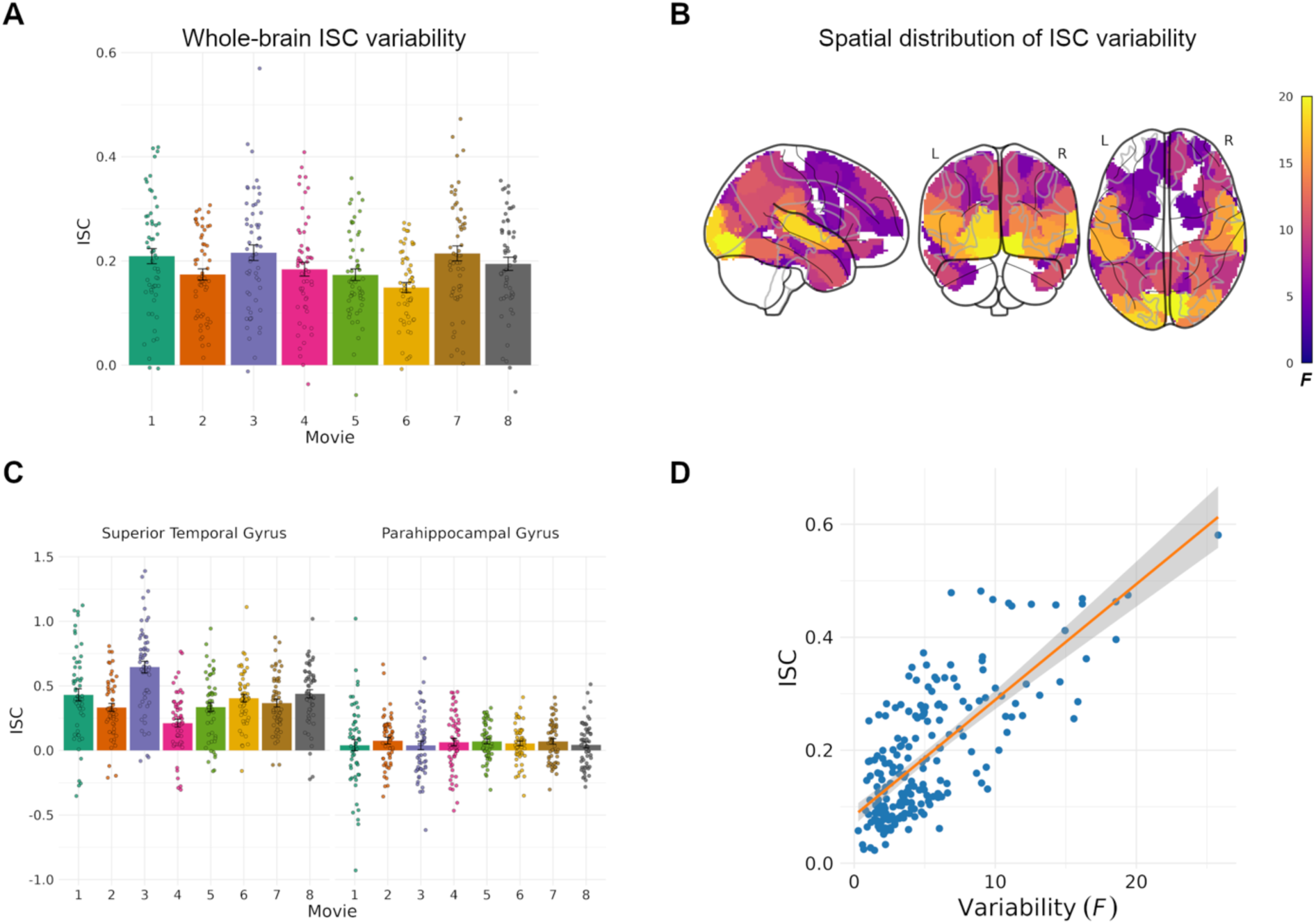
Variability in inter-subject correlations across movies in the CABB dataset. Fisher *z*-transformed ISC are shown for panels A, C, and D. **A)** Mean and standard error of whole-brain inter-subject correlation (ISC) values for each of the eight movies. Whole-brain ISC was computed by averaging ISC values across all cortical parcels for each subject and movie. A repeated-measures ANOVA revealed a significant main effect of Movie, indicating considerable variability in whole-brain ISC across movies. **B)** Spatial distribution of between-movie ISC variability across the brain. A significant main effect of Movie (family-wise error-adjusted *p* < 0.05) was found for 97 out of 210 cortical parcels (46.2%). The highest variability was observed in primary and associative visual cortices, as well as in primary and associative auditory cortices. Significant variability was also found in higher-order regions such as the precuneus and inferior parietal lobule. **C)** Brain regions with high between-movie variability, such as the superior temporal gyrus, tended to show relatively higher ISC levels, while regions with low variability, such as the parahippocampal gyrus, showed lower ISC. **D)** A strong positive correlation between ISC level (y-axis) and between-movie variability (x-axis) across brain regions indicates that regions with higher ISC levels also exhibit greater variability across movies. Each point represents one cortical parcel.

Next, we examined the spatial distribution of ISC variability across the cortex. For each of the 210 cortical parcels defined by the Brainnetome atlas, we conducted a repeated-measures ANOVA on ISC values with Movie as the within-subject factor. We found a statistically significant main effect of Movie in 97 of the 210 cortical parcels (46.2%; family-wise error [FWE]-adjusted *p* < 0.05). The strongest between-movie variability was observed in primary and associative visual cortex, as well as in primary and associative auditory cortex. Considerable variability was also observed in higher-order regions, including the precuneus and inferior parietal lobule. Parcel-wise statistical values quantifying between-movie variability are available online (https://dx.doi.org/10.17605/OSF.IO/S78WU). **Figure 3B** shows the spatial distribution of between-movie ISC variability across the cortex.

Importantly, these results were robust across additional control analyses. The pattern remained highly similar when using the Schaefer 300-parcel atlas (Schaefer et al., 2018) instead of the Brainnetome atlas (**Supplementary Figure 2**), when using equal-length movie segments to ensure that ISC differences were not solely driven by differences in movie duration (**Supplementary Figure 4**), and after residualizing ISC values for pairwise average FD, indicating that head motion did not explain between-movie ISC variability (**Supplementary Figure 6**). Results also did not depend on the specific choice of subject pairs, as shown by a resampling analysis in which the ANOVA was repeated for 1000 randomly sampled sets of non-overlapping subject pairs (**Supplementary Figure 7**).

To further test whether ISC patterns carry movie-specific information, we conducted a cross-validated multivariate classification analysis to distinguish movies based on parcel-wise ISC patterns. The classifier achieved an accuracy of 62.9%, substantially above chance level (12.5%; *p* = 0.001, permutation test). This indicates that the spatial distribution of ISC contains robust information about movie identity. The analysis is described in detail in the **Supplementary Results** and illustrated in **Supplementary Figure 8**.

### 3.2. Brain regions with stronger ISC show higher between-movie variability

We followed up on the finding of widespread between-movie ISC variability across the cortex by examining regions with particularly high or low variability. We discovered that brain regions showing high variability, such as the superior temporal cortex, also had relatively higher levels of ISC. Conversely, brain regions with low variability, such as the parahippocampal gyrus, had lower levels of ISC (**Figure 3C**). To quantify this relationship, we calculated the Pearson correlation between parcel-wise ISC level and between-movie ISC variability across the 210 cortical parcels. As shown in **Figure 3D**, ISC level was strongly positively correlated with between-movie variability (*r* = 0.73, *t*(208) = 15.62, *p* < 0.001, *R*^2^ = 0.54). This association remained strong after controlling for parcel-wise temporal signal-to-noise ratio calculated from preprocessed time series, partial *r* = .73, *t*(207) = 15.45, *p* < 0.001, *R*^2^ = 0.54 (**Supplementary Figure 9**).

The association between ISC level and ISC variability may partly arise because regions with stronger ISC provide a larger observable range in which between-movie differences can emerge. Accordingly, the positive correlation between ISC level and between-movie variability should not be taken as evidence for intrinsically more variable neural representations in visual and auditory cortex, but rather as evidence that stimulus-locked synchronization in these regions is particularly sensitive to differences between movies.

### 3.3. ISC–behavior associations vary across movies

Our next objective was to assess the consequences of between-movie ISC variability for the generalizability of ISC–behavior associations. We used IS-RSA to identify brain regions in which similarity between subjects in conceptual representations of novel objects, as measured by the *Features task*, was associated with ISC during movie watching. In other words, we tested whether subjects with more similar conceptual representations of the novel objects showed stronger neural synchronization in specific brain regions.

For each movie separately, we performed regression-based IS-RSA using linear mixed-effects models (G. Chen et al., 2017) to estimate the association between subject similarity in conceptual representations and ISC during movie watching, while controlling for similarities in lexical representations, assessed with a *Naming task*, as well as age and sex. Linear mixed-effects models are commonly used to quantify ISC–behavior associations (Baek et al., 2022, 2023; de Bruin et al., 2023; Finn et al., 2018; Jangraw et al., 2023; Moraczewski et al., 2018; van Baar et al., 2021; S. Ye et al., 2026). If between-movie ISC variability had little impact on ISC–behavior associations, we would expect similar IS-RSA patterns across movies. By contrast, if between-movie variability is consequential for ISC–behavior associations, we would expect IS-RSA patterns to differ across movies, and in the strongest case, to be movie-specific.

As shown in **Figure 4**, the IS-RSA revealed associations between conceptual representations of novel objects and ISC in distinct brain regions for each movie. For Movie #1, subject similarity in conceptual representations was significantly associated with neural synchronization in parcels within bilateral insula (right hemisphere: *p*_FWE_ = 0.004 and left hemisphere: *p*_FWE_ = 0.007), left inferior parietal lobule (*p*_FWE_ = 0.006), left precentral gyrus (*p*_FWE_ = 0.007), right superior frontal gyrus (*p*_FWE_ = 0.02), left inferior temporal gyrus (*p*_FWE_ = 0.03), and right anterior cingulate (*p*_FWE_ = 0.03). In contrast, for Movie #2, we found no statistically significant association between conceptual representations and neural synchronization. For Movie #3, significant associations were observed in the left precuneus (*p*_FWE_ < 0.05), with no overlap with the regions identified for Movie #1. For Movie #4, significant associations were observed in bilateral middle frontal gyrus (right hemisphere: *p*_FWE_ < 0.001 and left hemisphere: *p*_FWE_ = 0.03) and the right orbital gyrus (*p*_FWE_ = 0.02), again without overlap with the regions identified for Movies #1 or #3. Similarly, Movies #5 through #8 showed largely non-overlapping associations between conceptual representations and neural synchronization, with the exception of one parcel in the right middle temporal gyrus that was significant for both Movie #5 (*p*_FWE_ = 0.004) and Movie #6 (*p*_FWE_ < 0.001). Statistically significant IS-RSA associations for all movies are detailed in **Supplementary Table 2**. Full parcel-wise results for all movies are available online (https://dx.doi.org/10.17605/OSF.IO/S78WU).

**Figure 4.**
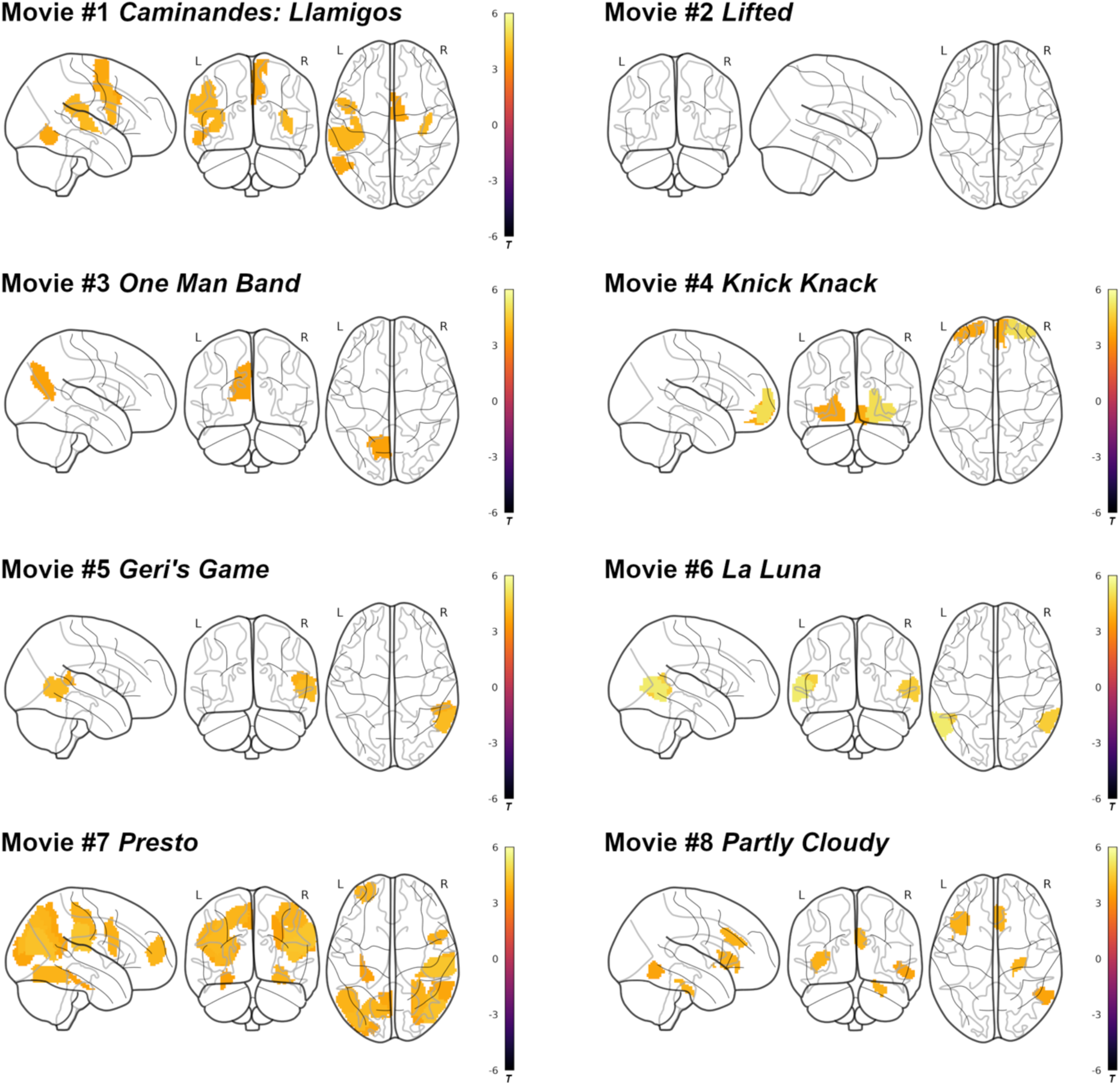
Between-movie ISC variability leads to varying IS-RSA associations between behavior and neural synchronization. We used inter-subject representational similarity analysis (IS-RSA) to identify brain regions where similarity in conceptual representations of novel objects was associated with similarity in neural synchronization during movie watching. The results revealed ISC–behavior associations in distinct sets of brain regions for each movie (family-wise error-adjusted *p* < 0.05).

Correlations between the movie-specific, unthresholded IS-RSA *t*-maps were low (mean *r* = 0.01 ± 0.13). This suggests limited consistency in the spatial patterns of ISC–behavior associations across movies, independent of the applied statistical threshold.

A control analysis using the Schaefer 300-parcel atlas instead of the Brainnetome atlas yielded similar results (**Supplementary Figure 3**). Because the higher number of parcels required stricter correction for multiple comparisons, fewer regions showed statistically significant IS-RSA effects, and Movie #8 showed no significant associations. In a further control analysis using equal-length movie segments, movie-specific IS-RSA patterns remained largely robust, although some effects changed after truncation (**Supplementary Figure 5**). The disappearance of the Movie #6 and #7 IS-RSA effects after truncating the time series further suggests that ISC–behavior associations may depend not only on the movie, but also on which segment of the movie is analyzed. Finally, a control analysis without covariates yielded highly similar IS-RSA results, indicating that the findings were robust and not driven by overlap with lexical representations or demographic similarity (**Supplementary Figure 10**).

### 3.4. A subset of IS-RSA effects generalize across movies

To examine whether any associations in the CABB dataset generalized across movies, we conducted an additional IS-RSA that included movie as a random effect. This analysis identified three brain regions, including bilateral posterior inferior temporal gyrus and right posterior superior temporal sulcus, where ISC was significantly associated with conceptual representations of novel objects across movies (**Supplementary Figure 11**). Because this model pooled data across the eight movies, these effects are best interpreted as average associations across the sampled movies with increased statistical precision, rather than as evidence that the model removes movie-specific ISC–behavior effects.

### 3.5. All main findings replicate across two additional open datasets

To assess the robustness of our findings across different subjects, movie types, and movie durations, we repeated all ISC analyses in two openly available movie-fMRI datasets: the HCP movie-watching dataset (Van Essen et al., 2013) and the Emo-FilM dataset (Morgenroth et al., 2025). In both datasets, we also conducted IS-RSA analyses using Big Five personality traits as a well-established behavioral construct.

#### 3.5.1. ISC varies across movies in HCP and Emo-FilM datasets

We first computed ISC across the cortex for each movie. As shown in **Supplementary Figures 12 and 13**, the overall spatial distribution of ISC evoked by each of the 28 movie clips and films in the two replication datasets largely resembled patterns reported in previous movie-fMRI studies and those observed in the CABB dataset. One notable exception was Movie #11 in the HCP dataset, which showed relatively low ISC outside visual cortex (X. Li et al., 2025).

Next, we tested whether whole-brain ISC varied between movies. In both replication datasets, the ANOVA revealed a significant main effect of Movie: HCP, *F*(13,1144) = 17.47, *p* < 0.001, η^2^_G_ = 0.13; Emo-FilM, *F*(13,169) = 2.59, *p* = 0.003, η^2^_G_ = 0.089. We then examined the spatial distribution of ISC variability across the cortex. For each of the 210 cortical parcels defined by the Brainnetome atlas, we conducted a repeated-measures ANOVA on ISC values with Movie as the within-subject factor. In both replication datasets, we found statistically significant between-movie ISC variability distributed across the cortex (FWE-adjusted *p* < 0.05). Parcel-wise statistical values quantifying between-movie variability are available online (https://dx.doi.org/10.17605/OSF.IO/S78WU). **Figure 5A** shows the spatial distribution of between-movie ISC variability in the HCP dataset, and **Figure 5B** shows the corresponding results for the Emo-FilM dataset.

**Figure 5.**
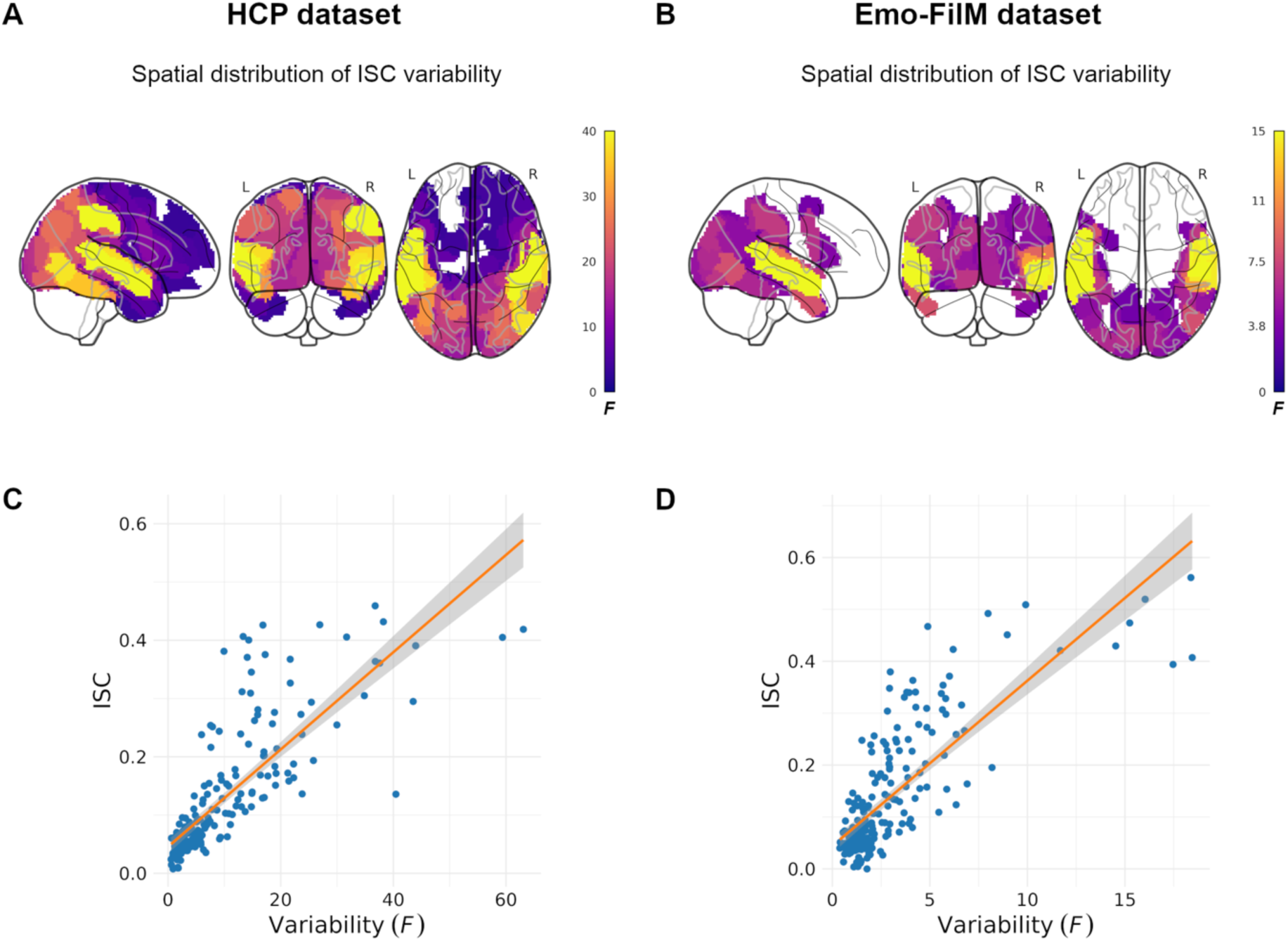
Variability in inter-subject correlations across movies in the HCP and Emo-FilM datasets. Fisher *z*-transformed ISC are shown for panels C and D. **A)** Spatial distribution of statistically significant between-movie ISC variability across the brain in the HCP dataset (family-wise error-adjusted *p* < 0.05). **B)** Spatial distribution of statistically significant between-movie ISC variability across the brain in the Emo-FilM dataset (family-wise error-adjusted *p* < 0.05). **C)** Strong positive correlation between ISC level (y-axis) and between-movie variability (x-axis) across brain regions in the HCP dataset. Each point represents one cortical parcel. **D)** Strong positive correlation between ISC level (y-axis) and between-movie variability (x-axis) across brain regions in the Emo-FilM dataset. Each point represents one cortical parcel.

For each replication dataset separately, we calculated the Pearson correlation between parcel-wise ISC level and between-movie ISC variability across the 210 cortical parcels. As shown in **Figures 5C and 5D**, ISC level was strongly positively correlated with between-movie variability: HCP, *r* = 0.80, *t*(208) = 19.16, *p* < 0.001, *R*^2^ = 0.64; Emo-FilM, *r* = 0.78, *t*(208) = 18.16, *p* < 0.001, *R*^2^ = 0.61. Together, these results show that our ISC findings were robust across independent datasets with live-action and animated movies of different durations.

#### 3.5.2. IS-RSA results for a well-validated construct vary across movies

We used IS-RSA to identify brain regions in which similarity between subjects in personality traits, measured with established questionnaires (John et al., 1991; McCrae & Costa, 2004), was associated with ISC during movie watching. In other words, we tested whether subjects with more similar personality profiles showed stronger neural synchronization in specific brain regions. These analyses conceptually extend prior work linking ISC to personality, in which associations were assessed across concatenated movie clips and therefore averaged across movie-specific effects (Finn et al., 2020; Matz et al., 2022).

For each replication dataset and movie separately, we performed regression-based IS-RSA using linear mixed-effects models. As shown in **Supplementary Figures 14 and 15**, IS-RSA revealed associations between personality similarity and ISC in distinct brain regions for each movie. Full parcel-wise results for all movies are available online (https://dx.doi.org/10.17605/OSF.IO/S78WU). Correlations between movie-specific, unthresholded IS-RSA t-maps were low in both replication datasets: HCP, mean *r* = 0.05 ± 0.13; Emo-FilM, mean *r* = 0.10 ± 0.15. This indicates limited consistency in the spatial pattern of ISC–behavior associations across movies, independent of the applied statistical threshold. Together, the IS-RSA results from the replication datasets show that movie-specific ISC–behavior associations are replicable across datasets and extend to a well-established behavioral construct.

## 4. Discussion

Here, we quantified the extent of ISC variability between different movies and assessed its consequences for ISC–behavior associations during movie watching. Across three independent datasets, our analyses revealed considerable between-movie variability in whole-brain ISC, and we found statistically significant variability distributed across the cortex. Between-movie ISC variability was present not only in early auditory and visual regions but also in higher-order regions, including areas of the default-mode network.

Using IS-RSA, we showed that associations between ISC and two distinct behavioral constructs had movie-specific spatial distributions, with minimal overlap between movies. In the following, we discuss the implications of these findings for the generalizability of ISC-based movie-fMRI results obtained from a single movie or a small set of movies.

### 4.1. What features of a movie drive neural synchronization?

A major challenge in using movies to investigate differences in neural synchronization is that it remains unclear which specific movie features drive ISC. Because resting-state fMRI time series are generally uncorrelated between subjects (Hasson et al., 2004), neural synchronization during movie watching must be driven by features of the movies. However, ever since the first studies using movie viewing as a paradigm in neuroscience (Bartels & Zeki, 2004; Hasson et al., 2004), movies have been recognized as highly complex, multidimensional sequences of stimuli, making it difficult to isolate the specific features that drive neural synchronization (Hasson et al., 2004, 2010). This challenge is further complicated by the fact that many movie features are correlated, making it difficult to isolate their individual contributions to neural synchronization (Rust & Movshon, 2005). Thus, even when movies are selected according to an experimenter-defined category, such as “Pixar-style animated short films,” individual films within this category will still vary along many dimensions that may differentially shape brain responses and give rise to distinct ISC patterns.

### 4.2. Characterizing movies in ISC-based analyses

Our findings suggest that different movies elicit distinct neural synchronization patterns across individuals, making them comparable to separate experimental tasks (Grall & Finn, 2022). Consequently, studies using movie paradigms should document stimulus properties with the same level of rigor applied in task-based research, even though the relevant features may be difficult to isolate (Vanderwal et al., 2019). Researchers should explicitly specify which perceptual, cognitive, or affective processes a given movie is intended to engage. Machine learning-based annotation tools, such as those implemented in *pliers* (McNamara et al., 2017) or *NeuroScout* (de la Vega et al., 2022), can help address this by automatically extracting auditory, visual, linguistic, or affective features from naturalistic materials. These tools provide a data-driven foundation for identifying features that may contribute to ISC and its variability across movies, offering a promising path toward greater interpretability and generalizability of ISC-based findings. However, current annotation tools are unlikely to fully capture higher-order narrative structure, social-affective nuance, or personal meaning, which may be particularly relevant for ISC variability in default-mode and other higher-order regions. Explaining variability in these regions will likely require complementing automated feature extraction with human ratings or theory-driven narrative annotations (Schmälzle & Huskey, 2023). Recent work also highlights the potential of combining stimulus-based annotations with multi-dimensional experience sampling, which captures subjects’ ongoing thought patterns without disrupting the movie-watching experience (Wallace et al., 2025). Combining such subjective reports with automated feature annotations could clarify how both stimulus features and internal cognitive states shape ISC.

### 4.3. Selecting movies for a target construct

Prior ISC-based movie-fMRI studies have sometimes selected theoretically motivated movies to target a specific construct of interest, such as political advertisements or debates to examine differences in political orientation (Katabi et al., 2023; van Baar et al., 2021) or emotionally evocative scenes to study depressive symptoms (Guo et al., 2015). In such cases, the link between the selected movie and the behavioral construct is central to the interpretation of ISC effects. In the CABB dataset, the animated short films were selected to include object categories that were frequently associated with the novel objects in pilot testing, such as plants, toys, and food. However, we did not quantify the occurrence of the specific feature dimensions assessed in the Features task. Accordingly, some movie-specific IS-RSA effects may partly reflect differences in construct relevance: certain movies may have provided more opportunities than others for conceptual representations to shape shared neural responses.

This issue is unlikely to be specific to our study. As discussed above, even movies selected from the same broad category can differ in many features that may affect ISC. The difficulty of selecting an appropriate movie also depends on the construct of interest. For emotional responses, relevant stimulus properties such as valence or arousal can be quantified relatively directly (Morgenroth et al., 2023). By contrast, it is less clear what would make a movie optimal for broad constructs such as personality traits or clinical diagnoses (Eickhoff et al., 2020). This is particularly relevant for attempts to use ISC as a biomarker or stable trait marker: Our findings suggest strongly synchronized regions may be highly sensitive to the movie selected. Candidate ISC-based markers should therefore be evaluated across multiple theoretically relevant movies. Beyond theory-driven stimulus selection, a complementary approach is to treat movie selection itself as an empirical question (X. Li et al., 2025).

### 4.4. Including the stimulus as a random effect in the statistical model

A potential solution to the limited generalizability of ISC-based movie-fMRI findings is to include movie as a random effect in the statistical model. Following Yarkoni (2022), “random effects are used to model […] variables that are assumed to be stochastically sampled from some underlying population and can vary across replications without meaningfully altering the research question.” (Yarkoni, 2022 p. 3). With rare exceptions (van Baar et al., 2021), studies investigating group differences in ISC or ISC–behavior associations across multiple movies have typically treated stimulus as fixed, either by modeling movie as a fixed effect, estimating separate models for each movie, or concatenating movie time series before fitting a single model. This limits the ability to generalize beyond the specific stimulus set used, a problem long recognized in neuroscience (Westfall et al., 2017) and other fields (Clark, 1973; Coleman, 1964; Yarkoni, 2022). The phenomenon we describe here may therefore reflect a specific instance of the “stimulus-as-fixed-effect fallacy”, which, in principle, can be addressed by including movie as a random effect. To explore this approach in the CABB dataset, we tested an IS-RSA model that included movie as a random effect. This analysis revealed consistent associations in a small number of brain regions, but most effects remained movie specific. However, because our random-effects model included only eight movies, variance components for the movie effect were estimated from a small number of movies, limiting the precision of these estimates and the strength of conclusions about generalization beyond the sampled movies (Judd et al., 2012).

Including movie as a random effect leaves open the question of the broader stimulus population to which the results are intended to generalize. In a random-effects framework, the included movies are treated as samples from a population of possible movies that could have been used to address the same research question (Yarkoni, 2022). For this assumption to be meaningful, that population needs to be defined. In many controlled task-fMRI settings, the relevant stimulus population can be specified relatively clearly. For example, an auditory experiment may sample pure tones of different frequencies, with the aim of generalizing to other frequencies that were not explicitly tested. In this case, the stimulus dimension of interest is explicit and can be systematically sampled. By contrast, the relevant stimulus population is less straightforward for movies, which differ along many perceptual, narrative, social, and affective dimensions. The intended population could be animated short films, live-action social narratives, emotionally negative movie clips, or movies containing specific social interactions. These choices matter because different movie features may contribute to ISC and ISC–behavior associations. Thus, modeling movie as a random effect can account statistically for variability across the included movies, but it does not identify which movie features drive this variability or determine how far beyond the sampled movies the results should generalize.

### 4.5. Constraints on generality

It remains an open question to what extent our findings generalize across different uses of dynamic, multidimensional materials in neuroscience. For example, studies employing control conditions such as scrambled movies (Hasson, Yang, et al., 2008; Lerner et al., 2011) or scrambled music (Abrams et al., 2013) may be less affected by the stimulus-specificity in ISC observed here. The rationale behind scrambled conditions is that many (though likely not all) features of the stimulus materials that drive variability in ISC are held constant between the intact and scrambled versions, isolating only those of interest to the researcher. However, if this assumption is not fully met, for example because it is difficult to match all features that variably drive ISC between the real and control conditions, stimulus-specificity could still pose a challenge. Our findings may also be less consequential for studies focusing on specific types of events embedded within dynamic, multidimensional materials, such as those investigating memory or event segmentation (Baldassano et al., 2017; Ben-Yakov & Henson, 2018; J. Chen et al., 2017), many of which use the general linear model to examine brain responses to specific event types (Bartels & Zeki, 2004; Jacoby et al., 2016; Richardson, 2019; Richardson et al., 2018). In such cases, variability between entire movies may have minimal impact on the targeted event-level effects. Our findings are not relevant for studies using voxel-wise encoding models to map neural responses to annotated stimulus features, as these models are typically evaluated on independent datasets, providing a direct assessment of generalizability across different stimuli (Dupré la Tour et al., 2025; Gifford et al., 2025; Huth et al., 2016).

Within-subject functional connectivity measures may be more consistent across different movies than ISC-based measures (Tian et al., 2021). This relative stability is likely because functional connectivity is largely driven by intrinsic network architecture (Buckner et al., 2013), which is commonly measured during rest and shows only modest modulation by movies (Vanderwal et al., 2015) or conventional task conditions (Cole et al., 2014; Gratton et al., 2018). However, movie-specific differences in functional connectivity may still be meaningful in certain contexts (Kröll et al., 2023; Vanderwal et al., 2017), such as when using functional connectivity patterns to predict individual phenotypes (Finn & Bandettini, 2021).

More broadly, the implications of ISC variability for the generalizability of ISC–behavior associations are unlikely to be limited to movie fMRI. They may extend to other dynamic, multidimensional stimulus materials, such as spoken narratives, audiobooks, or music; to analytical approaches such as inter-subject functional correlation (M. Ye et al., 2023) and ISC-based predictive modeling of individual phenotypes (X. Li et al., 2025); and to neuroimaging modalities beyond fMRI, including magnetoencephalography (MEG), electroencephalography (EEG) (Petroni et al., 2018), or functional near-infrared spectroscopy (fNIRS). Most of these possibilities remain to be tested and require systematic empirical investigation in future research.

### 4.6. Limitations and future directions

A limitation of our study is that all three datasets used relatively short films or movie clips. It therefore remains to be determined whether ISC patterns and ISC–behavior associations show comparable movie-specificity for feature-length movies, for example from the naturalistic neuroimaging database (Aliko et al., 2020).

An important future direction concerns the role of functional alignment. Our analyses were based on anatomically aligned, parcel-averaged time series and therefore did not account for individual differences in fine-grained functional topographies. Methods such as hyperalignment or the shared-response model address this issue by estimating subject-specific transformations into a common functional space and increase sensitivity to shared responses across individuals (P.-H. (Cameron) Chen et al., 2015; Guntupalli et al., 2016; Haxby et al., 2011, 2020). Future studies analyzing voxel- or vertex-level responses to feature-length movies could use such data to estimate robust functional-alignment transformations. Alternatively, datasets with several sufficiently long movie clips could use leave-one-movie-out cross-validation, training the alignment on all but one movie clip and testing whether movie-specific ISC and ISC–behavior associations persist for the held-out clip.

Future work could also compare variability in responses to movies with variability across cognitive tasks that target similar underlying processes (Andermane et al., 2019; Cai et al., 2024; Fedorenko et al., 2013). Such comparisons may help contextualize movie-induced variability relative to more traditional task-based paradigms. Making such a comparison meaningful would require identifying the specific perceptual, cognitive, or affective processes that a given set of movies is likely to engage.

### 4.7. Conclusion

Across three independent datasets, we found that ISC patterns vary across movies. This between-movie variability was reflected in distinct spatial distributions of ISC–behavior associations, with limited overlap between movies. These findings suggest that ISC–behavior associations can be strongly movie-specific, underscoring the need to link conclusions to detailed characterizations of the movies used, particularly when findings are based on a single movie or a small set of movies.

## Data and Code availability

Raw data from the CABB dataset are available in the Radboud Data Repository, as described in the accompanying data descriptor (https://doi.org/10.1016/j.neuroimage.2022.119734). The HCP data are publicly available and can be accessed through the HCP website (https://db.humanconnectome.org/). The Emo-FilM dataset is publicly available on OpenNeuro (https://openneuro.org/datasets/ds004892). The *Python* and *R* code used for the analyses is available on Github (https://github.com/simonleipold/movie_variability) and archived on Zenodo (https://doi.org/10.5281/zenodo.13348505).

## Author Contributions

**Simon Leipold:** Conceptualization, Formal analysis, Writing – Original Draft, Writing – Review & Editing, Visualization, Funding acquisition. **Rajat Ravi Rao:** Methodology, Formal analysis, Writing – Review & Editing. **Jan-Mathijs Schoffelen:** Formal analysis, Writing – Review & Editing. **Sara Bögels:** Investigation, Project administration, Writing – Review & Editing. **Ivan Toni:** Conceptualization, Writing – Original Draft, Writing – Review & Editing, Supervision, Project administration, Funding acquisition.

## Acknowledgments

This work was supported by the Dutch Research Council (NWO; Gravitation Grant 024.001.006 to the Language in Interaction Consortium), by the Swiss National Science Foundation (SNSF; Grants 206557 and 222073 to Simon Leipold), and the European Research Council (ERC; Advanced Grant 101054559 to Ivan Toni).

The HCP movie-watching dataset was provided by the Human Connectome Project, WU-Minn Consortium (Principal Investigators: David Van Essen and Kamil Ugurbil; 1U54MH091657) funded by the 16 NIH Institutes and Centers that support the NIH Blueprint for Neuroscience Research; and by the McDonnell Center for Systems Neuroscience at Washington University.

Acquisition of the Emo-FilM dataset was funded by the Swiss National Science Foundation (SNSF; Grant 180319 to Patrik Vuilleumier, Ronan Boulic, and Dimitri Van de Ville) and benefited from resources of the Swiss Center of Affective Sciences at UNIGE. It was also supported by Grant #2021-613 of the Strategic Focus Area “Personalized Health and Related Technologies (PHRT)” of the ETH Domain (Swiss Federal Institutes of Technology).

