## Supplementary for "Inter-subject correlations and their behavioral associations vary across movies: Implications for generalizability"

### Supplementary Methods

#### 1. *CABB dataset*

##### *1.1. Movie stimuli*

###### *1.1.1. Caminandes: Llamigos*

*Description:* Computer-animated slapstick short film set in a snowy Patagonian landscape. The film depicts a llama and a penguin competing over food in a visually simple, fast-paced sequence.

*Target audience:* General/family audience
*Producer:* Ton Roosendaal
*Director:* Pablo Vazquez
*Year released:* 2016
*Length:* 2 min 14 sec

###### *1.1.2. Lifted*

*Description:* Pixar animated short film in which a young alien attempts to abduct a sleeping human under the supervision of an instructor.

*Target audience:* General/family audience
*Producer:* Katherine Sarafian
*Director:* Gary Rydstrom
*Year released:* 2006
*Length:* 4 min 21 sec

###### *1.1.3. One Man Band*

*Description:* Pixar animated musical-comedy short film in which two street musicians compete for the attention and coin of a young girl.

*Target audience:* General/family audience
*Producer:* Osnat Shurer
*Director:* Andrew Jimenez and Mark Andrews
*Year released:* 2005
*Length:* 3 min 37 sec

###### *1.1.4. Knick Knack*

*Description:* Pixar animated short film about a snowman toy trapped inside a snow globe who repeatedly tries to escape.

*Target audience:* General/family audience
*Producer:* John Lasseter
*Director:* John Lasseter
*Year released:* 1989
*Length:* 2 min 53 sec

###### *1.1.5. Geri's Game*

*Description:* Pixar animated short film in which an older man plays chess against himself in a park.

*Target audience:* General/family audience
*Producer:* Karen Dufilho
*Director:* Jan Pinkava
*Year released:* 1997
*Length:* 4 min 05 sec

###### *1.1.6. La Luna*

*Description:* Pixar animated short film about a boy who joins his father and grandfather on a nocturnal journey by boat and discovers their unusual work on the moon.

*Target audience:* General/family audience
*Producer:* Kevin Reher
*Director:* Enrico Casarosa
*Year released:* 2011
*Length:* 6 min 05 sec

###### *1.1.7. Presto*

*Description:* Pixar animated short film in which a magician's performance is disrupted by his hungry rabbit.

*Target audience:* General/family audience
*Producer:* Richard Hollander
*Director:* Doug Sweetland
*Year released:* 2008

*Length:* 4 min 25 sec

###### *1.1.8. Partly Cloudy*

*Description:* Pixar animated short film about clouds that create babies and storks that deliver them.

*Target audience:* General/family audience
*Producer:* Kevin Reher
*Director:* Peter Sohn
*Year released:* 2009
*Length:* 5 min 00 sec

##### *1.2. fMRIprep preprocessing pipeline*

Results included in this manuscript come from preprocessing performed using *fMRIPrep* 20.2.7 (Esteban et al., 2019, 2022) (RRID:SCR_016216), which is based on *Nipype* 1.7.0 (Esteban et al., 2021; Gorgolewski et al., 2011) (RRID:SCR_002502).

###### *1.2.1. Anatomical data preprocessing*

The T1-weighted (T1w) image was corrected for intensity non-uniformity (INU) with *N4BiasFieldCorrection* (Tustison et al., 2010), distributed with *ANTs* 2.3.3 (Avants et al., 2008) (RRID:SCR_004757), and used as T1w-reference throughout the workflow. The T1w-reference was then skull-stripped with a *Nipype* implementation of the *antsBrainExtraction.sh* workflow (from *ANTs*), using *OASIS30ANTs* as target template. Brain tissue segmentation of cerebrospinal fluid (CSF), white-matter (WM) and gray-matter (GM) was performed on the brain-extracted T1w using *fast* (*FSL* 5.0.9, RRID:SCR_002823) (Zhang et al., 2001). Brain surfaces were reconstructed using *recon-all* (*FreeSurfer* 6.0.1, RRID:SCR_001847) (Dale et al., 1999), and the brain mask estimated previously was refined with a custom variation of the method to reconcile *ANTs*-derived and *FreeSurfer*-derived segmentations of the cortical gray-matter of *Mindboggle* (RRID:SCR_002438) (Klein et al., 2017). Volume-based spatial normalization to one standard space (*MNI152NLin2009cAsym*) was performed through nonlinear registration with *antsRegistration* (*ANTs* 2.3.3), using brain-extracted versions of both T1w reference and the T1w template. The following template was selected for spatial normalization: *ICBM 152 Nonlinear Asymmetrical template version 2009c* (Fonov et al., 2009) [RRID:SCR_008796; TemplateFlow ID: *MNI152NLin2009cAsym*].

###### *1.2.2. Functional data preprocessing*

For the one BOLD run found per subject (across all tasks and sessions), the following preprocessing was performed. First, a reference volume and its skull-stripped version were generated by aligning and averaging one single-band reference (SBRefs). A B0-nonuniformity map (or fieldmap) was estimated based on two (or more) echo-planar imaging (EPI) references with opposing phase-encoding directions, with *3dQwarp* (Cox & Hyde, 1997) (*AFNI* 20160207). Based on the estimated susceptibility distortion, a corrected EPI (echo-planar imaging) reference was calculated for a more accurate co-registration with the anatomical reference. The BOLD reference was then co-registered to the T1w reference using *bbregister* (*FreeSurfer*) which implements boundary-based registration (Greve & Fischl, 2009). Co-registration was configured with six degrees of freedom. Head-motion parameters with respect to the BOLD reference (transformation matrices, and six corresponding rotation and translation parameters) are estimated before any spatiotemporal filtering using *mcflirt* (*FSL* 5.0.9) (Jenkinson et al., 2002). BOLD runs were slice-time corrected to 0.445 s (0.5 of slice acquisition range 0 s-0.89 s) using *3dTshift* from *AFNI* 20160207 (Cox & Hyde, 1997) (RRID:SCR_005927). First, a reference volume and its skull-stripped version were generated using a custom methodology of *fMRIPrep*. The BOLD time series (including slice-timing correction when applied) were resampled onto their original, native space by applying a single, composite transform to correct for head-motion and susceptibility distortions. These resampled BOLD time series will be referred to as preprocessed BOLD in original space, or just preprocessed BOLD. The BOLD time series were resampled into standard space, generating a preprocessed BOLD run in *MNI152NLin2009cAsym* space. First, a reference volume and its skull-stripped version were generated using a custom methodology of *fMRIPrep*. Several confounding time series were calculated based on the preprocessed BOLD: framewise displacement (FD), DVARS and three region-wise global signals. FD was computed using two formulations following Power (absolute sum of relative motions) (Power et al., 2014) and Jenkinson (relative root mean square displacement between affines) (Jenkinson et al., 2002). FD and DVARS are calculated for each functional run, both using their implementations in *Nipype* (following the definitions by Power et al. 2014). The three global signals are extracted within the CSF, the WM, and the whole-brain masks. The head-motion estimates calculated in the correction step were also placed within the corresponding confounds file. The confound time series derived from head motion estimates and global signals were expanded with the inclusion of temporal derivatives and quadratic terms for each (Satterthwaite et al., 2013). Frames that exceeded a threshold of 0.5 mm FD or 1.5 standardized DVARS were annotated as motion outliers. All resamplings can be performed with a single interpolation step by composing all the pertinent transformations (i.e. head-motion transform matrices, susceptibility distortion correction when available, and co-registrations to anatomical and output spaces). Gridded (volumetric) resamplings were performed using *antsApplyTransforms* (*ANTs*), configured with Lanczos interpolation to minimize the smoothing effects of other kernels (Lanczos, 1964). Non-gridded (surface) resamplings were performed using *mri_vol2surf* (*FreeSurfer*).

Many internal operations of *fMRIPrep* use *Nilearn* 0.6.2 (Abraham et al., 2014) (RRID:SCR_001362), mostly within the functional processing workflow. For more details of the pipeline, see the section corresponding to workflows in *fMRIPrep*'s documentation.

##### *1.3. Inter-subject representational similarity analysis*

We used inter-subject representational similarity analysis (IS-RSA) to identify brain regions where subject similarity in conceptual representations of novel objects, as estimated in the Features task, was associated with neural synchronization between subjects during movie watching. Below, we describe the stimuli and behavioral tasks used to estimate conceptual and lexical representations of novel objects, the generation of representational dissimilarity matrices (RDM), and provide details on the regression-based IS-RSA. This analysis linked conceptual representations to neural synchronization during movie watching, while controlling for lexical representations of the novel objects, as well as similarities in sex and age. Portions of these descriptions are adapted from Eijk et al. (2022). Note that the Communicative Alignment of Brain and Behaviour (CABB) dataset encompasses neural and behavioral data for both a pre-session and a post-session. Here, we exclusively analyzed data from the post-session because the movie-watching fMRI data was acquired during the post-session.

###### *1.3.1. Stimuli*

The stimuli consisted of 16 images of novel objects, which were composite geometric figures known as "Fribbles" (see **Figure 1C**), adapted from Barry et al. (2014). These objects were colored blue and displayed on a light gray background. The adaptations from the original stimuli were guided by pilot tests aimed at ensuring that each Fribble would evoke different conceptualizations across individuals. The Fribbles consisted of various appendages attached to the same core element, a cup-like shape.

###### *1.3.2. Behavioral tasks*

Two different behavioral tasks, the *Features task* and the *Naming task*, were performed individually by each subject in dedicated soundproof cubicles while sitting in front of a 24-inch screen (BENQ 24" Full HD [1920 x 1080] x 120 Hz LED TFT), delivering their responses with a keyboard and a mouse.

In the *Features task*, each Fribble was presented in the top left corner of the screen, next to the statement, "To what extent do you view this picture as …". Subjects were asked to use a linear visual analog scale to report how well each Fribble matched 29 different features (e.g., pointy, symmetrical, human, related to movement; full list in **Supplementary Table 1**) (Binder et al., 2016). The left end of the scale (0) represented no match (or: not applicable), while the right end of the scale (100) represented a very strong match between the Fribble on the screen and the indicated feature. For each feature, subjects were asked to report their judgment within a few seconds by moving a red cursor (initially placed in the middle of the scale) along a white scale using the mouse. Only after subjects had moved the cursor for each of the 29 features could they proceed to the next trial.

In the *Naming task*, the 16 Fribbles were presented together on the screen, with their positions randomized across subjects. On each trial, subjects were asked to name the Fribble framed in a red square using one to three words of their choice, which would help the other member of the pair identify that Fribble among the others displayed on the screen. The arrangement of the Fribbles on the screen, as well as the order in which the to-be-named Fribbles were presented, was randomized across subjects.

###### *1.3.3. Conceptual representational dissimilarity matrix*

In the *Features task*, subjects were asked to report on linear visual analog scales how well each Fribble matched 29 different features. The position of the red cursor on the white analog scale was transformed into a centesimal score, resulting in a vector of 29 feature scores per Fribble for each subject. **Supplementary Figure 1** visualizes the distribution of Feature ratings for each feature and Fribble. Distances between each pair of vectors of 29 feature scores for a Fribble were calculated using the Mahalanobis distance. This was done both within each subject and across subjects, resulting in a large conceptual representational dissimilarity matrix (RDM).

###### *1.3.4. Lexical representational dissimilarity matrix*

In the *Naming task*, subjects were asked to write down three words of their choice to name each of the Fribbles. The text provided by subjects underwent a standardization process that involved removing specific characters (indicated here between <>: <‘>, <“>, <()>, <&>, <+>, <.>, <;>), converting the characters <-> and </> into spaces, transforming <=> into the word <is>, correcting clear spelling mistakes, eliminating uppercase letters, and converting numerals into their word equivalents. Using *SNAUT* (Mandera et al., 2017), these words were then cross-referenced with a corpus used for *word2vec* analysis (Mikolov, Chen, et al., 2013; Mikolov, Sutskever, et al., 2013). Words not found in the corpus were corrected where feasible, and compounds not listed were split into separate words. Cosine distances between each pair of names on the list were calculated with *SNAUT*, resulting in a large lexical RDM. Cosine distances for each pair of names were determined based on the frequency of co-occurrence within the NLPL Dutch CoNLL17 corpus (Zeman et al., 2017).

###### *1.3.5. Inter-subject representational dissimilarity matrices*

To assess the average dissimilarity of subjects’ conceptual and lexical representations of the Fribbles, we reduced the large conceptual and lexical RDMs to subject-by-subject RDMs (112 x 112) that served as input for the IS-RSA. In detail, for both the large conceptual and lexical RDMs, we first removed all within-subject distances from the matrix. Second, we removed all distances between separate stimuli from the matrix (e.g., distances between the first and second Fribble stimulus). This resulted in a matrix that exclusively included distances between different subjects but only for the same stimuli. This step ensured that we did not compare different subjects' representations of different stimuli. Third, we averaged the distances across the 16 stimuli for each possible pair of subjects, resulting in a subject-by-subject matrix (112 x 112), representing the average distance of subjects' conceptual and lexical representations of the novel objects. These subject-by-subject RDMs served as input for the IS-RSA, along with the ISC matrix for each movie.

###### *1.3.6. Inter-subject correlation distance matrices*

For the movie-watching fMRI data, we re-calculated the inter-subject correlation (ISC) matrices for each parcel, which represent neural synchronization in that particular brain region between all possible subject pairs, as ISC *distance* matrices, reflecting the dissimilarities in fMRI time series, using *scikit-learn* (version 1.0.2; RRID:SCR_002577). This step ensured that positive associations reflected a positive link between conceptual representations and neural synchronization (i.e. greater subject similarity would be associated with stronger neural synchronization). The correlation distance was calculated as 1 minus the Pearson correlation between the parcel-wise fMRI time series. Note that this step only changes the sign of the association, not its strength or statistical significance.

###### *1.3.7. Linear mixed-effects models*

We conducted a regression-based IS-RSA (Finn et al., 2020; Nummenmaa et al., 2012) using linear mixed-effects models (Chen et al., 2017; van Baar et al., 2021) to link subject similarity in conceptual representations of the novel objects with neural synchronization during movie watching. The linear mixed-effects models were estimated using the *lme4* package (version 1.1-33; RRID:SCR_015654) in *R* (version 4.1.0; RRID:SCR_001905). Model assumptions were checked using visualizations from the *performance* package (version 0.10.2) (Lüdecke et al., 2021). We estimated a full model to simultaneously assess the main effects of subjects' similarities in conceptual representations on their neural synchronization during movie watching, while controlling for subject similarities in lexical representations of the novel objects, sex, and age.

In detail, for each movie and for each cortical parcel of the Brainnetome atlas (Fan et al., 2016), we estimated a linear mixed-effects model using vectorized and *z*-transformed full subject-by-subject RDMs (upper and lower triangles, excluding the diagonal) as input. Because the full RDMs include data duplication, we adjusted the degrees of freedom as described in Chen et al. (2017). The full model included the ISC distances during the viewing of that particular movie between all possible pairs of subjects (i,j) as the dependent variable, modeled by the following:

(1) a main effect of pairwise distance in conceptual representations,
(2) a main effect of pairwise distance in lexical representations,
(3) a main effect of pairwise difference in sex (same, different),

(4) a main effect of pairwise difference in age,
(5) a random intercept for subject [i] within the pair,
(6) a random intercept for subject [j] within the pair.

The model also included a group intercept which is not explicitly coded in *lme4*.

Statistical inference was performed on the beta coefficients of the main effect of pairwise distance in conceptual representations of the novel objects on neural synchronization during movie watching, using a one-sample *t*-test against zero. The degrees of freedom were adjusted to 6211, which equals the number of all possible subject pairs (6216) minus the number of fixed effects in the model (5; intercept and four main effects) (Baek et al., 2022; Chen et al., 2017). Tests of main effects were performed one-sided to investigate positive associations between similarities in conceptual representations and neural synchronization. The statistical significance level was set to α = 0.05, family-wise error (FWE)-adjusted for the number of parcels.

#### 2. HCP dataset

##### 2.1. Movie stimuli

###### 2.1.1. Two Men

*Description:* A man sees another man running past him on a road in rural Australia and reflects on the runner's possible motives. The film contains accented English dialogue and subtitles.

*Target audience:* Adults
*Producer:* Dominic Allen
*Director:* Dominic Allen
*Year released:* 2009
*Length:* 4 min 05 sec

###### 2.1.2. Welcome to Bridgeville

*Description:* Documentary-style clip in which people describe why they value living in a small American town, interleaved with scenes from community life.

*Target audience:* General/family audience
*Producer:* Jill Silberstein

*Director:* Phillip Van
*Year released:* 2011
*Length:* 3 min 41 sec

###### *2.1.3. Pockets*

*Description:* Close-up documentary short showing people holding objects they keep in their pockets and explaining the personal significance of these items.

*Target audience:* General/family audience
*Producer:* Andrew Hinton
*Director:* James Lees
*Year released:* 2008
*Length:* 3 min 08 sec

###### *2.1.4. Inside the Human Body*

*Description:* Short inspirational montage showing people who have overcome physical disabilities.

*Target audience:* General/family audience
*Producer:* N/A
*Director:* N/A
*Year released:* 2011
*Length:* 1 min 03 sec

###### *2.1.5. Inception*

*Description:* Excerpt from Inception in which two characters explore a dream world and learn about its unusual physical and emotional properties.

*Target audience:* Adults
*Producer:* Emma Thomas and Christopher Nolan
*Director:* Christopher Nolan
*Year released:* 2010
*Length:* 3 min 48 sec

###### *2.1.6. The Social Network*

*Description:* Excerpt from The Social Network depicting Mark Zuckerberg's disciplinary hearing at Harvard and its aftermath in a fictionalized portrayal.

*Target audience:* Adults
*Producer:* Scott Rudin, Dana Brunetti, Michael De Luca, and Ceán Chaffin
*Director:* David Fincher
*Year released:* 2010
*Length:* 4 min 19 sec

###### ***2.1.7. Ocean's Eleven***

*Description:* Excerpt from Ocean's Eleven in which Danny Ocean and his accomplices meet to plan a Las Vegas casino heist.

*Target audience:* Adults
*Producer:* Jerry Weintraub
*Director:* Steven Soderbergh
*Year released:* 2001
*Length:* 4 min 10 sec

###### ***2.1.8. Off the Shelf***

*Description:* A small red flower escapes from its pot and travels through a neighborhood. The clip follows a simple visually driven journey.

*Target audience:* General/family audience
*Producer:* N/A
*Director:* N/A
*Year released:* 2007
*Length:* 3 min 00 sec

#### ***2.1.9. 1212***

*Description:* A man and a woman have a metaphysical encounter in a hotel room.

*Target audience:* Adults
*Producer:* N/A
*Director:* N/A
*Year released:* N/A
*Length:* 3 min 05 sec

###### *2.1.10. Mrs. Meyer's Clean Day*

*Description:* Documentary-style clip about an urban vegetable garden serving community needs.

*Target audience:* General/family audience
*Producer:* N/A
*Director:* N/A
*Year released:* N/A
*Length:* 3 min 23 sec

###### *2.1.11. Northwest Passage*

*Description:* Montage of dreary landscapes and abandoned structures set to eerie music.

*Target audience:* Adults
*Producer:* N/A
*Director:* N/A
*Year released:* N/A
*Length:* 2 min 23 sec

###### *2.1.12. Home Alone*

*Description:* Excerpt from Home Alone in which the main character walks through the house and realizes that he is alone.

*Target audience:* General/family audience
*Producer:* John Hughes
*Director:* Chris Columbus
*Year released:* 1990
*Length:* 3 min 54 sec

###### *2.1.13. Erin Brockovich*

*Description:* Excerpt from Erin Brockovich in which the main character meets with a plaintiff and then visits a legal office with her young children.

*Target audience:* Adults
*Producer:* Danny DeVito, Michael Shamberg, and Stacey Sher
*Director:* Steven Soderbergh
*Year released:* 2000

*Length:* 3 min 51 sec

###### *2.1.14. The Empire Strikes Back*

*Description:* Excerpt from The Empire Strikes Back showing a scene from a rebel base on an icy planet, including an argument between Leia and Han Solo.

*Target audience:* Adults
*Producer:* Gary Kurtz
*Director:* Irvin Kershner
*Year released:* 1980
*Length:* 4 min 15 sec

#### *3. Emo-FilM dataset*

##### *3.1. Movie stimuli*

###### *3.1.1. After The Rain*

*Description:* A man contemplates the meaning of life and human interaction on a rainy day in the city. The film is an existential drama centered on daily routine, doubt, and altered perception.

*Target audience:* Adults
*Producer:* N/A
*Directors:* Marco Lucà and Laura Aloi
*Year released:* 2012
*Length:* 8 min 16 sec

###### *3.1.2. Between Viewings*

*Description:* A disillusioned estate agent is asked to sell his childhood home and is forced to re-examine his life. The film combines comedy and drama around memory, personal history, and self-reflection.

*Target audience:* Adults
*Producer:* Vaia Ikonomou
*Director:* Raphaël Biss
*Year released:* 2011

*Length:* 13 min 28 sec

###### ***3.1.3. Big Buck Bunny***

*Description:* A large, gentle rabbit is harassed by a group of bullying forest animals and eventually retaliates. The film is a computer-animated comedy with exaggerated slapstick action and animal characters.

*Target audience:* General/family audience
*Producer:* Ton Roosendaal
*Director:* Sacha Goedegebure
*Year released:* 2008
*Length:* 8 min 10 sec

###### ***3.1.4. Chatter***

*Description:* A girl witnesses a disturbing event online; after the electricity is cut off inside her apartment and later returns, she feels that she is not alone. The film is a short thriller built around isolation, suspense, and threat.

*Target audience:* Adults
*Producer:* Leo Resnes
*Director:* Leo Resnes
*Year released:* 2010
*Length:* 6 min 45 sec

###### ***3.1.5. First Bite***

*Description:* A teenage girl discovers the power of seduction. The film is categorized as a romance and focuses on interpersonal attraction and emerging sexuality.

*Target audience:* Adults
*Producer:* Thomas Done
*Director:* Louise Marie Cooke
*Year released:* 2011
*Length:* 9 min 59 sec

###### ***3.1.6. Lesson Learned***

*Description:* A young man turns his life around after becoming involved in gang violence. The film is a drama about peer influence, violence, and the consequences of risky social environments.

*Target audience:* Adults
*Producer:* Fritz Joseph
*Director:* Fritz Joseph
*Year released:* 2012
*Length:* 11 min 07 sec

###### ***3.1.7. Payload***

*Description:* A man must sacrifice everything to save his family. The film is a science-fiction drama set in a dystopian future, combining family conflict, survival, and social inequality.

*Target audience:* Adults
*Producer:* Thomas Bicknell
*Director:* Stuart Willis
*Year released:* 2012
*Length:* 16 min 48 sec

###### ***3.1.8. Sintel***

*Description:* A young woman embarks on a dangerous quest to find her lost friend, a dragon. The film is an animated fantasy story involving attachment, loss, danger, and discovery.

*Target audience:* Adults
*Producer:* Ton Roosendaal
*Director:* Colin Levy
*Year released:* 2010
*Length:* 12 min 02 sec

###### ***3.1.9. Spaceman***

*Description:* A young man sets out on a curious and unusual path to realize his dream of becoming an astronaut. The film is a romance/drama with a science-fiction framing and an emphasis on aspiration, eccentricity, and interpersonal longing.

*Target audience:* Adults
*Producer:* N/A
*Director:* Jono Schaferkotter
*Year released:* 2009
*Length:* 13 min 25 sec

###### *3.1.10. Superhero*

*Description:* A single mother cares for her son with terminal cancer. The film is an emotionally heavy drama about illness, caregiving, courage, and parental love.

*Target audience:* Adults
*Producer:* Langley McArol and Dino Muccio
*Director:* Langley McArol
*Year released:* 2011
*Length:* 17 min 08 sec

###### *3.1.11. Tears of Steel*

*Description:* A group of soldiers and scientists attempt to stop an army of robots threatening the planet by correcting a past mistake. The film combines live action and visual effects in an action/science-fiction narrative.

*Target audience:* Adults
*Producer:* Ton Roosendaal
*Director:* Ian Hubert
*Year released:* 2012

*Length:* 9 min 48 sec

###### *3.1.12. The Secret Number*

*Description:* A psychiatrist is compelled by his patient, an obsessive mathematician, to consider the existence of a secret integer between three and four. The film is a psychological drama with science-fiction elements and philosophical themes.

*Target audience:* Adults
*Producer:* Roque Nonini and Frank Ponce
*Director:* Colin Levy
*Year released:* 2012
*Length:* 13 min 04 sec

###### *3.1.13. To Claire; From Sonny*

*Description:* A young man writes a letter to his first true love. The film is a short drama centered on memory, affection, and romantic reflection.

*Target audience:* Adults
*Producer:* N/A
*Director:* Josh Beattie
*Year released:* 2010
*Length:* 6 min 42 sec

###### ***3.1.14. You Again***

*Description:* A chance encounter between two former high-school sweethearts forces them to confront the ways they have, and have not, changed. The film is a romance/drama about memory, identity, and reconnection.

*Target audience:* Adults
*Producer:* N/A
*Director:* N/A
*Year released:* N/A
*Length:* 13 min 18 sec

### Supplementary Results

#### Assessing robustness of findings using a different cortical parcellation

To evaluate whether our findings were robust to atlas choice, we repeated all ISC and IS-RSA analyses in the CABB dataset using time series extracted from the Schaefer 300-parcel atlas (Schaefer et al., 2018) instead of the Brainnetome atlas (Fan et al., 2016).

##### *Between-movie ISC variability*

We first repeated the parcel-wise ANOVAs using time series extracted from the Schaefer atlas. A significant main effect of Movie (FWE-adjusted $p < 0.05$) was observed in 137 of 300 cortical parcels (45.7%), confirming that distributed between-movie ISC variability was robust to atlas choice (**Supplementary Figure 2A**). Parcel-wise statistical values quantifying between-movie variability in ISC are available online (https://dx.doi.org/10.17605/OSF.IO/S78WU).

We also repeated the analysis examining the relationship between ISC level and between-movie variability. The Pearson correlation between average ISC and between-movie variability across the 300 parcels confirmed a strong positive association ($r = 0.68$, $t(298) = 16.14$, $p < 0.001$, $R^2 = 0.47$) (**Supplementary Figure 2B**).

##### *IS-RSA*

We repeated the regression-based IS-RSA using time series extracted from the Schaefer atlas for all movies. As in the main analysis, we used linear mixed-effects models to relate pairwise similarity in subjects' conceptual representations of novel objects to ISC, while controlling for similarity in lexical representations, age, and sex. The results again showed movie-specific associations between conceptual similarity and ISC in distinct brain regions, with minimal overlap across movies (**Supplementary Figure 3**). Full parcel-wise results for all movies are available online (https://dx.doi.org/10.17605/OSF.IO/S78WU).

#### Controlling for differences in movie length using truncated BOLD time series

To evaluate whether differences in movie duration influenced our findings in the CABB dataset, we conducted a control analysis in which all ISC and IS-RSA analyses were repeated using truncated time series of equal length. Specifically, we limited each movie's BOLD time series to the first 2 minutes and 14 seconds, matching the duration of the shortest stimulus in the CABB dataset (Movie #1). This approach ensured that our main results were not confounded by unequal numbers of time points across movies. All other preprocessing and analysis steps remained identical to those described in the main analyses.

##### *Variability in ISC between movies of equal length*

First, we re-ran the repeated-measures ANOVA on Fisher *z*-transformed ISC values. As in the main analysis, whole-brain ISC values were computed by averaging ISC across all cortical parcels for each subject pair and movie. A repeated-measures ANOVA on these values (with non-overlapping subject pairs) again revealed a significant main effect of Movie ($F(7,385) = 8.38$, $p < 0.001$, $\eta^2_G = 0.097$), indicating robust variability in ISC across movies even when movie duration is held constant. **Supplementary Figure 4A** visualizes the means and standard errors of the whole-brain ISC values for each movie separately.

We also repeated the parcel-wise ANOVAs using the truncated time series. A significant main effect of Movie (FWE-adjusted $p < 0.05$) was observed in 93 of 210 cortical parcels (44.3%), confirming that between-movie ISC variability is not attributable to unequal movie lengths (**Supplementary Figure 4B**).

We also re-ran the analysis examining the relationship between ISC level and between-movie variability using equal-length time series. As in the main analysis, brain regions with higher ISC tended to show greater variability across movies. A Pearson correlation between average ISC and between-movie variability across the 210 parcels confirmed this association ($r = 0.55$, $t(208) = 9.48$, $p < 0.001$, $R^2 = 0.30$), indicating that this relationship remains robust when controlling for movie duration (**Supplementary Figure 4C and 4D**).

##### *IS-RSA using movies of equal length*

We repeated the regression-based IS-RSA using truncated time series of equal length across all movies. As in the main analysis, we used linear mixed-effects models to relate pairwise similarity in subjects' conceptual representations of novel objects to ISC, while controlling for similarity in lexical representations, age, and sex. The results again showed distinct brain regions associated with conceptual similarity for each movie, with minimal overlap across movies (**Supplementary Figure 5**). This confirms that the movie-specific IS-RSA patterns observed in the main analysis were not driven by differences in movie duration.

#### Controlling for head motion quantified by framewise displacement (FD)

In the CABB dataset, overall head motion was low, with mean FD below 0.2 mm for all movies. However, FD differed significantly between movies, as shown by a repeated-measures ANOVA with Movie as the within-subject factor, $F(7, 777) = 11.88$, $p < 0.001$. Mean FD showed a gradual increase across the scan, suggesting an order effect whereby subjects moved their heads more over time (mean ± standard deviation): Movie #1 = 0.137 ± 0.076 mm, Movie #2 = 0.150 ± 0.060 mm, Movie #3 = 0.151 ± 0.046 mm, Movie #4 = 0.160 ± 0.054 mm, Movie #5 = 0.157 ± 0.045 mm, Movie #6 = 0.171 ± 0.090 mm, Movie #7 = 0.171 ± 0.053 mm, and Movie #8 = 0.174 ± 0.056 mm.

Because mean FD differed across movies, we repeated the ISC variability analysis after controlling for head motion. For each subject and movie, FD was averaged across the corresponding movie segment from the *fMRIprep* confound file. For each subject pair and movie, pairwise FD was then defined as the average FD of the two subjects in that pair. For each parcel separately, Fisher $z$-transformed ISC values were residualized with respect to pairwise FD using a linear model. The parcel mean was added back to the residuals to preserve the original scale for visualization. These FD-residualized ISC values were then entered into the same repeated-measures ANOVA framework as in the main analysis, with Movie as the within-subject factor and subject pair as the repeated-measures unit. As shown in **Supplementary Figure 6**, the results remained virtually unchanged, confirming that the observed between-movie ISC variability was not explained by differences in head motion between movies.

#### Resampling subject pairs to test the robustness of the ISC repeated-measures ANOVA

In our main analysis quantifying between-movie variability in the CABB dataset, we used a subset of 56 non-overlapping subject pairs to ensure statistical independence required for repeated-measures ANOVA. To assess whether our results were robust to the specific choice of subject pairs, we conducted a resampling analysis. Specifically, we randomly sampled 1000 sets of 56 non-overlapping subject pairs from the full ISC matrix and repeated the ANOVA on whole-brain ISC values for each sample. This allowed us to estimate the stability of the $F$-statistic across valid pair configurations. The mean $F$-statistic across iterations was 3.88, with 98% of samples exceeding the critical value for $\alpha = 0.05$ ($F = 2.03$). The originally reported $F$-statistic ($F = 4.93$) fell well within this distribution (**Supplementary Figure 7)**. These results confirm that our findings are not driven by a specific subject-pairing scheme and are robust across different valid pair selections.

#### Multivariate classification of movies based on ISC patterns

To determine whether spatial patterns of ISC values robustly differentiate between movies in the CABB dataset, we implemented a multivariate classification analysis using elastic net-regularized multinomial logistic regression. Analyses were conducted in *R* using the *caret* (version 7.0-1; RRID:SCR_021138) package in combination with *glmnet* (version 4.1-8; RRID:SCR_015505).

To ensure independence of samples, we used the same 56 subject pairs as in the ANOVA described in the main text, such that each subject contributed to a single ISC value per movie. ISC values were reshaped into a feature matrix where each row represented a unique subject pair-movie combination, and each column corresponded to a cortical parcel. Feature matrices were standardized (mean-centered and scaled to unit variance) prior to classification.

A five-fold cross-validation procedure was used to train the classifier, with regularization via elastic net as implemented in *glmnet*. Out-of-fold predictions from cross-validation were extracted and compared to the true movie labels to compute a cross-validated confusion matrix and overall classification accuracy.

The classification analysis yielded an overall cross-validated accuracy of 62.9% (chance level = 12.5% for eight movies). Using a permutation test (1000 iterations with shuffled movie labels), we confirmed that the observed classification accuracy of 62.9% was significantly above chance ($p$ = 0.001), indicating that the spatial distribution of ISC across the cortex contains robust, discriminative information about movie identity. **Supplementary Figure 8** displays the corresponding confusion matrix and permutation-based null distribution.

#### Controlling for parcel-wise temporal signal-to-noise ratio

Brain regions with higher signal quality may yield more reliable ISC estimates, which could potentially contribute to the association between ISC level and between-movie variability. In the CABB dataset, we therefore tested whether this association was explained by regional differences in temporal signal-to-noise ratio (tSNR). tSNR was calculated from voxel-wise preprocessed time series and averaged across subjects and movies for each parcel. We then computed a partial Pearson correlation between ISC level and between-movie ISC variability across all parcels, controlling for parcel-wise tSNR using the *ppcor* (version 1.1) package in *R*. As shown in **Supplementary Figure 9**, the association remained strong after controlling for tSNR (partial $r$ = .73, $p$ < .001). This indicates that the relationship between ISC level and between-movie variability was not explained by regional differences in tSNR.

#### Omitting covariates in the IS-RSA model

To evaluate the robustness of our IS-RSA findings in the CABB dataset, we re-ran the analyses without including the covariates (i.e., Naming RDM, age similarity, and sex similarity). For each movie separately, we estimated the association between ISC and pairwise similarity in conceptual representations from the *Features task* using linear mixed-effects models, identical to those in the main analysis but excluding the covariates. The overall pattern of results remained highly similar to the original models (**Supplementary Figure 10**), confirming that the observed associations between conceptual similarity and neural synchronization were not driven by overlap with lexical representations or demographic similarity.

#### Incorporating a random effect for movie in the IS-RSA

To assess the impact of modeling between-movie variability in the IS-RSA in the CABB dataset, we conducted an additional regression-based analysis that included a random intercept for movie. This analysis used the same full linear mixed-effects model described above for the individual movie analyses but was fit to the data from all movies simultaneously, with movie treated as a random effect. The procedure for model estimation and parcel-wise statistical inference was identical to that used in the separate models for each movie. Note that our stimulus set of eight movies may not be sufficiently large to support robust random-effects modeling or strong generalization across movies (Judd et al., 2012).

As shown in **Supplementary Figure 11**, the IS-RSA model incorporating a random intercept for movie revealed significant associations between the conceptual representations of the novel objects and ISC in three brain regions: bilateral posterior inferior temporal gyrus and right posterior superior temporal sulcus (**Supplementary Table 3**). Notably, these effects showed partial overlap with the results from the movie-specific models. The left inferior temporal parcel was also statistically significant for Movie #1, but not for any other movie. Similarly, the right inferior temporal parcel showed a significant association with ISC only in Movie #8, and the right posterior superior temporal sulcus parcel was significant only in Movie #6. These findings suggest that although some regions exhibit consistent associations with conceptual representations across movies, most regions appear to be movie specific.

### Supplementary Figures

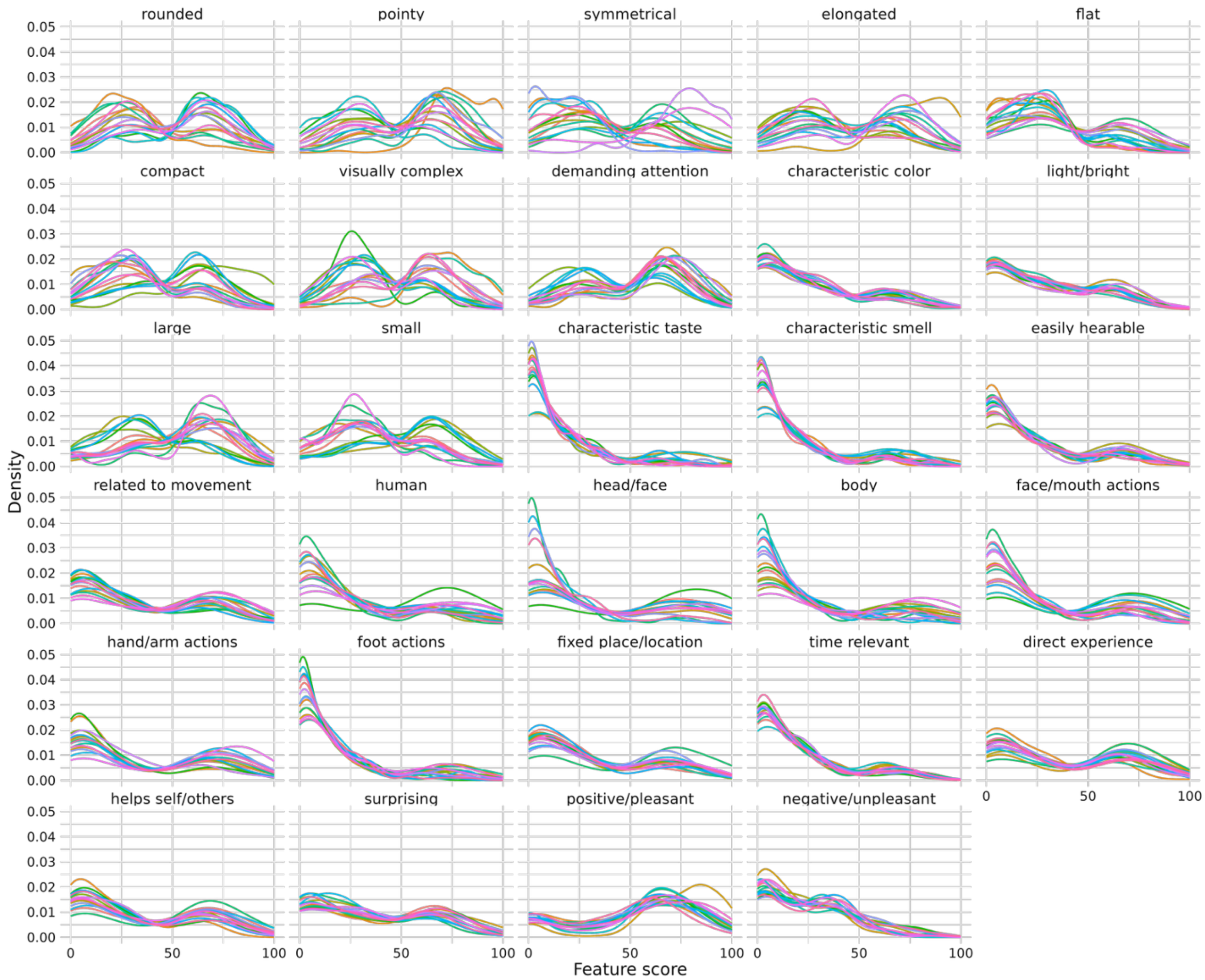


**Supplementary Figure 1. Distribution of Feature ratings in the *Features task*.** Density plots show the distribution of ratings for each of the 29 features. Each panel represents one feature, and differently colored lines indicate the distributions for the 16 Fribbles. Ratings were given on a visual analog scale ranging from 0 to 100, with higher values indicating a stronger perceived match between the Fribble and the respective feature.

#### Control analysis: Schaefer atlas

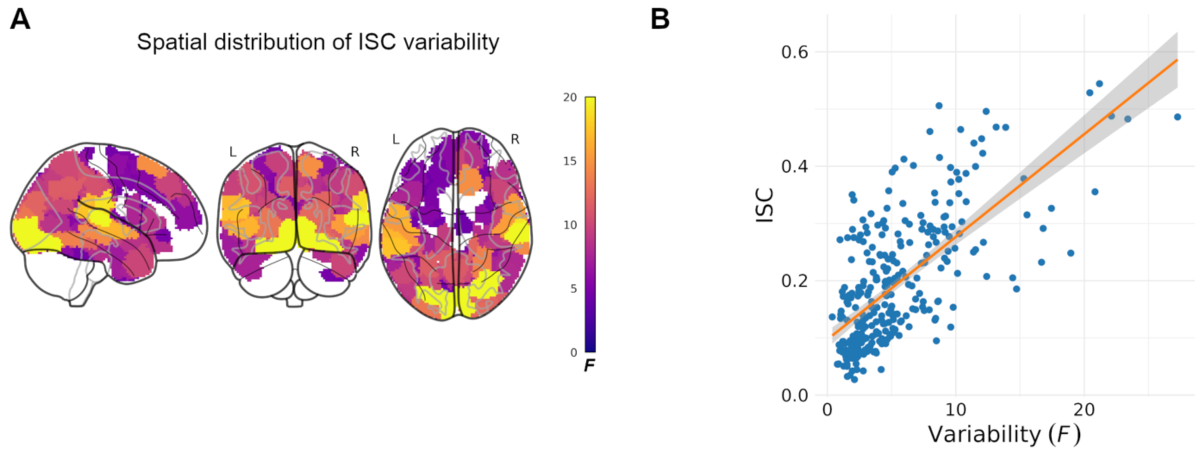


**Supplementary Figure 2. Between-movie variability in ISC in CABB dataset using Schaefer 300-parcel atlas.** Fisher *z*-transformed ISC are shown for panel B. **A)** Spatial distribution of statistically significant between-movie ISC variability across the brain in the CABB dataset (family-wise error-adjusted $p < 0.05$). **B)** Strong positive correlation between ISC level (y-axis) and between-movie variability (x-axis) across brain regions in the CABB dataset. Each point represents one cortical parcel of the Schaefer 300-parcel atlas (Schaefer et al., 2018).

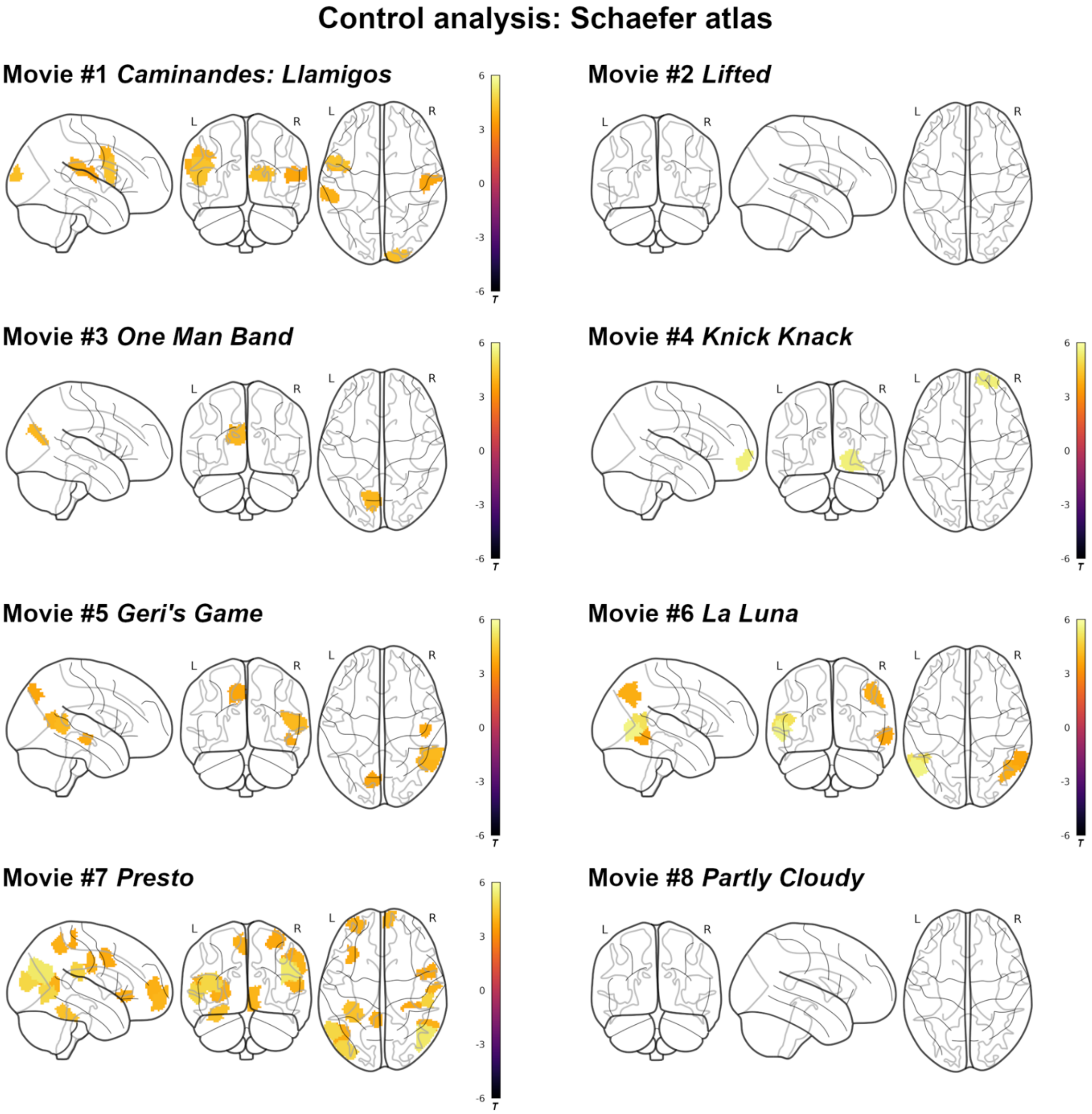


**Supplementary Figure 3. IS-RSA in CABB dataset using Schaefer 300-parcel atlas.**

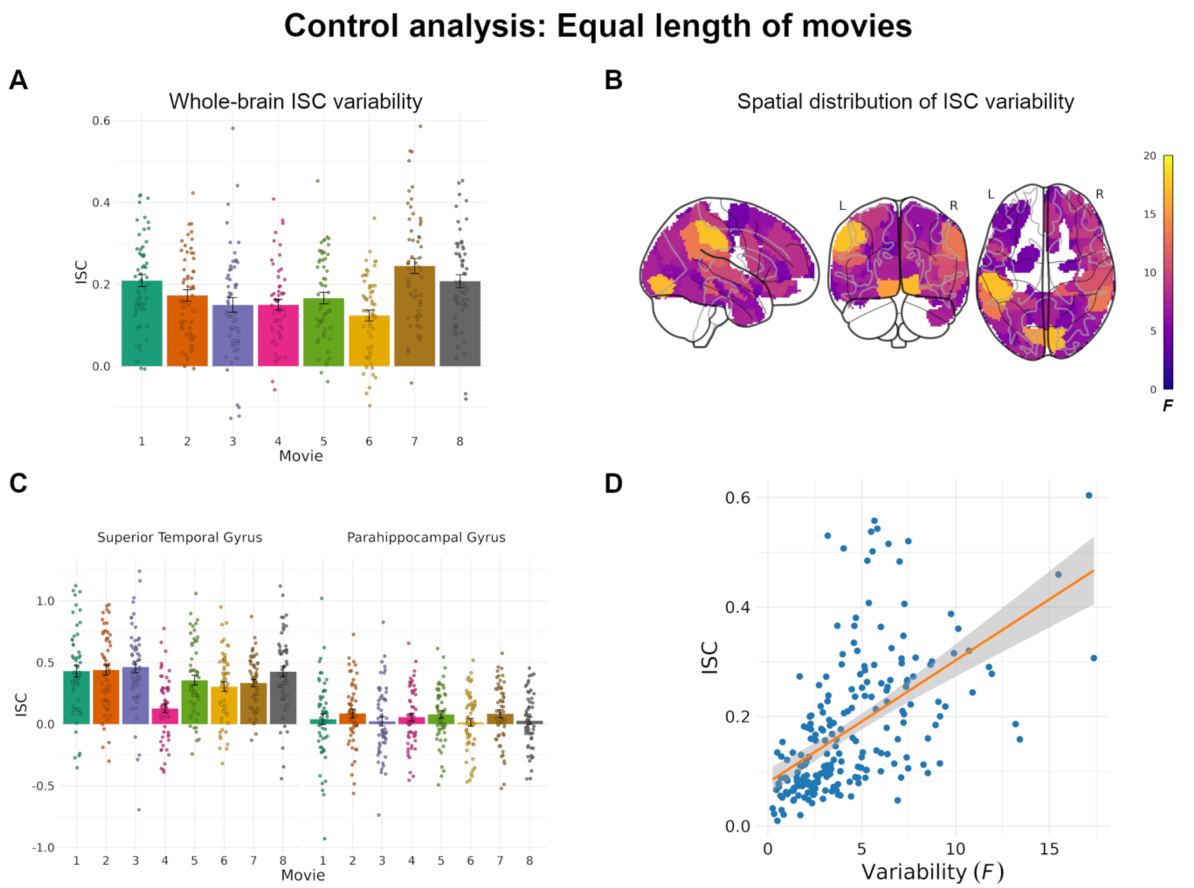


**Supplementary Figure 4. Between-movie variability in ISC in CABB dataset using equal-length movie segments.** Fisher *z*-transformed ISC are shown for panels A, C, and D. **A)** Mean and standard error of whole-brain inter-subject correlation (ISC) values for each of the eight movies. **B)** Spatial distribution of between-movie ISC variability across the brain (family-wise error-adjusted $p < 0.05$). **C)** Brain regions with high between-movie variability, such as the superior temporal gyrus, tended to show relatively higher ISC levels, while regions with low variability, such as the parahippocampal gyrus, showed lower ISC. **D)** A strong positive correlation between ISC level (y-axis) and between-movie variability (x-axis) across brain regions indicates that regions with higher ISC levels also exhibit greater variability across movies. Each point represents one cortical parcel of the Brainnetome atlas (Fan et al., 2016).

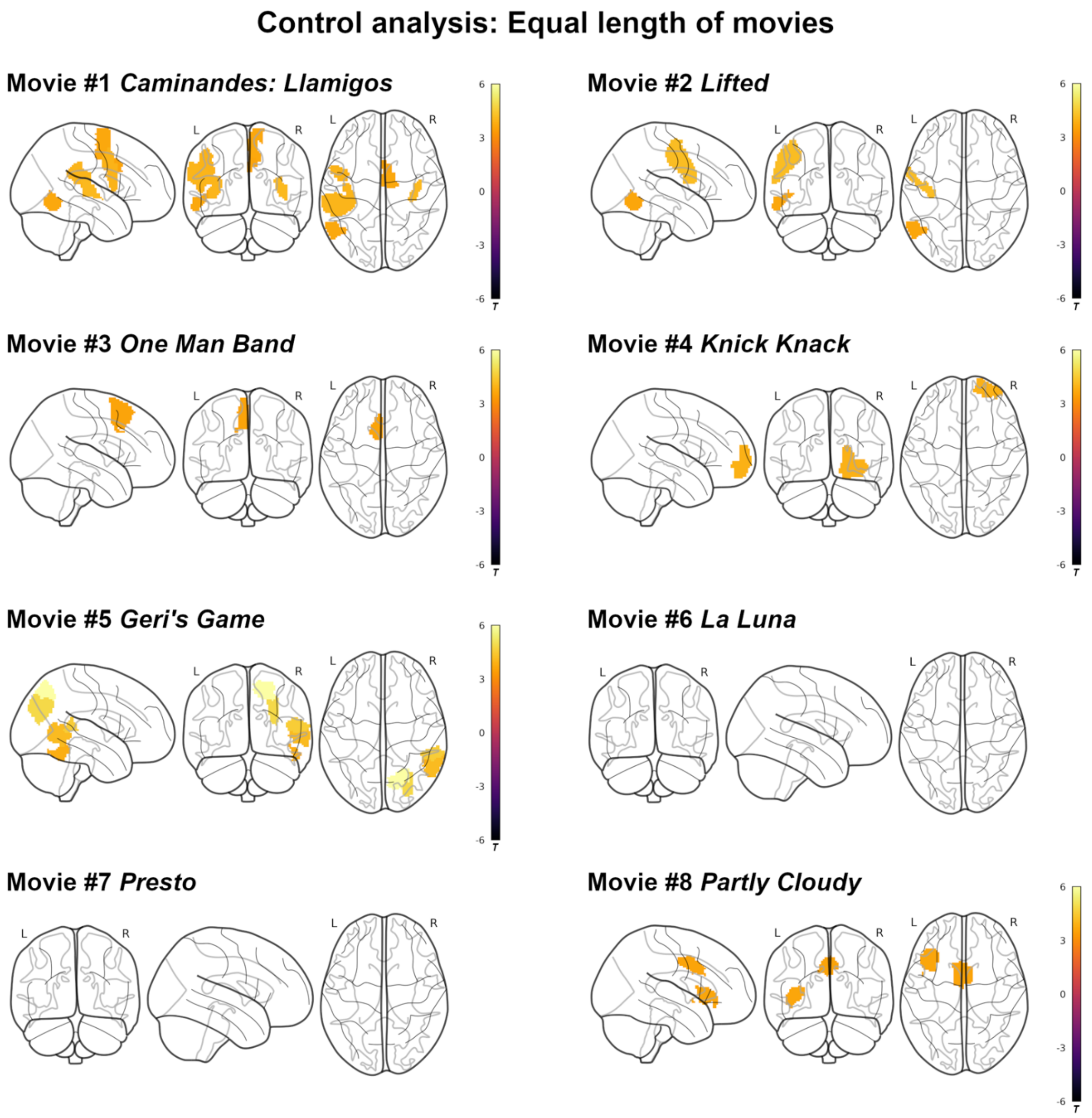


**Supplementary Figure 5. IS-RSA in CABB dataset using equal-length movie segments.** Associations between pairwise similarity in conceptual representations of novel objects and ISC values for each movie, based on truncated BOLD time series of equal duration.

#### Control analysis: Head motion

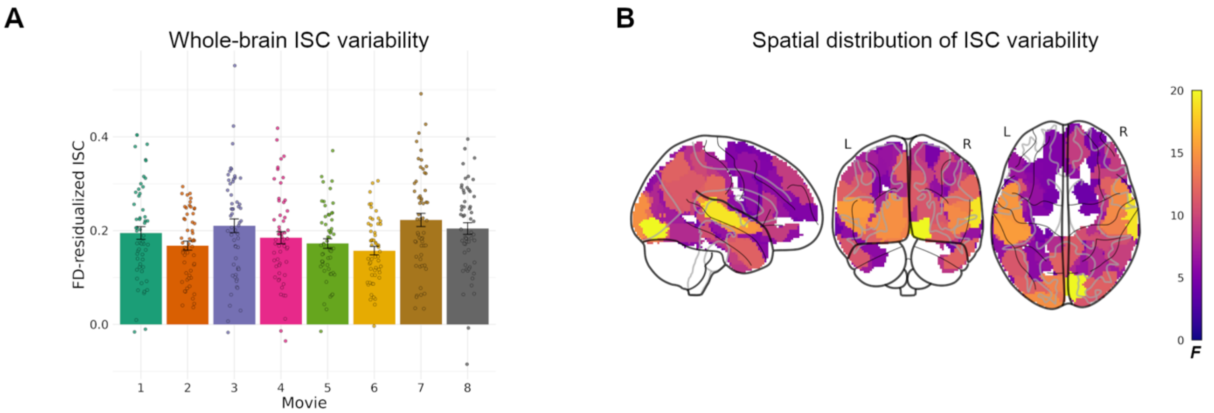


**Supplementary Figure 6. Head motion control analysis in the CABB dataset.** Between-movie ISC variability remained robust after controlling for head motion. ISC values were residualized with respect to pairwise average framewise displacement (FD) before repeating the main repeated-measures ANOVA. **A)** Mean and standard error of FD-residualized whole-brain ISC values for each movie. **B)** Spatial distribution of between-movie variability in FD-residualized ISC across cortical parcels.

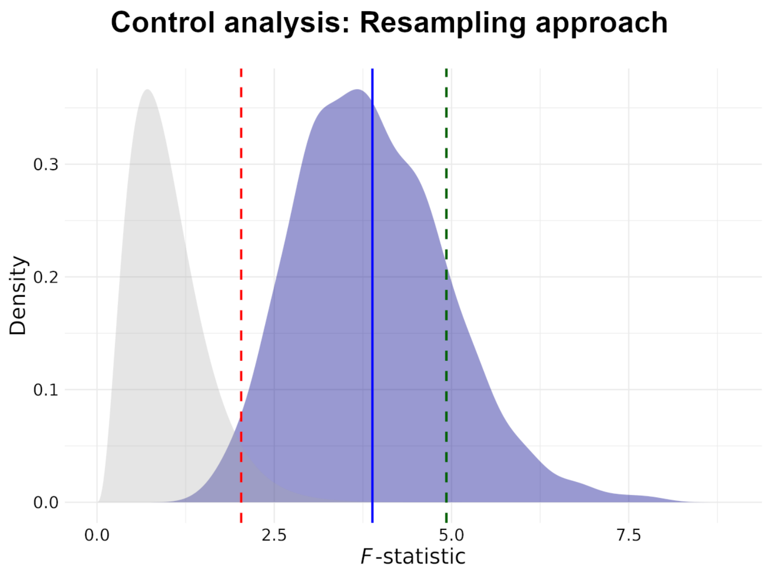


**Supplementary Figure 7. Resampling-based control analysis.** The blue distribution shows the empirical density of *F*-statistics obtained from 1000 repeated-measures ANOVAs on randomly sampled, non-overlapping subject pairs. Each ANOVA tested for differences in whole-brain ISC across movies. The dashed red line marks the theoretical *F*-critical value ($\alpha$ = 0.05); the dashed green line marks the *F*-statistic from the original analysis based on a fixed set of 56 non-overlapping pairs.

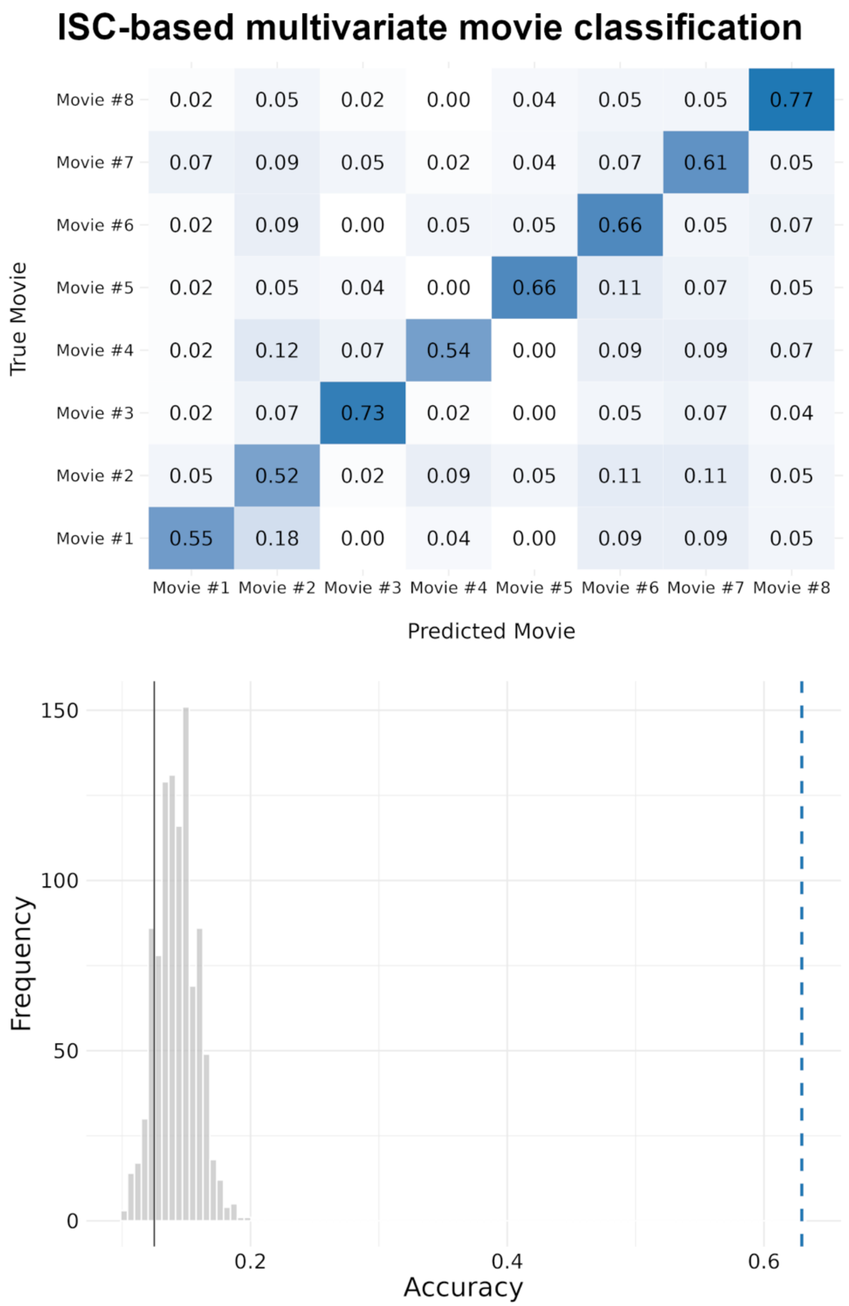


**Supplementary Figure 8. ISC-based multivariate movie classification.** Confusion matrix (top) and permutation-based distribution of classification accuracy (bottom). The dashed blue line marks the empirically observed accuracy, and the solid gray line indicates chance level.

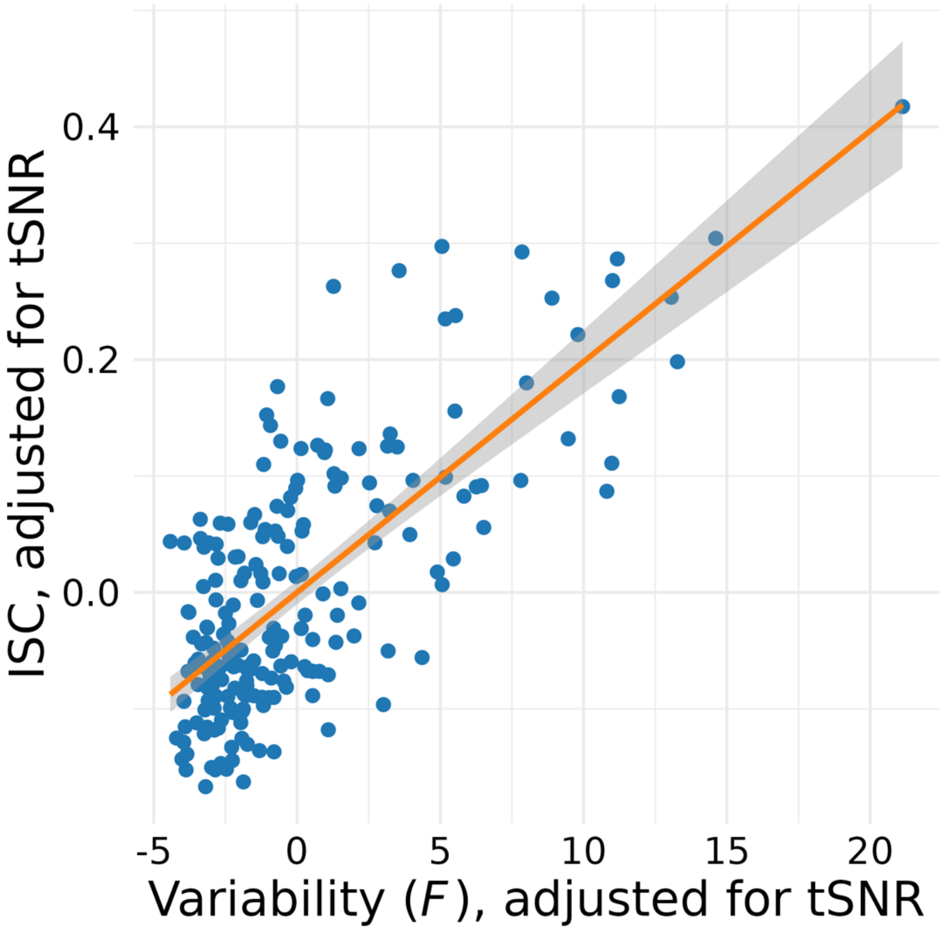


**Supplementary Figure 9. ISC level–variability association after controlling for temporal signal-to-noise ratio in the CABB dataset.** The association between parcel-wise ISC level and between-movie ISC variability remained strong after controlling for parcel-wise temporal signal-to-noise ratio (tSNR). Both axes show residualized values after regressing out tSNR. Each point represents one cortical parcel.

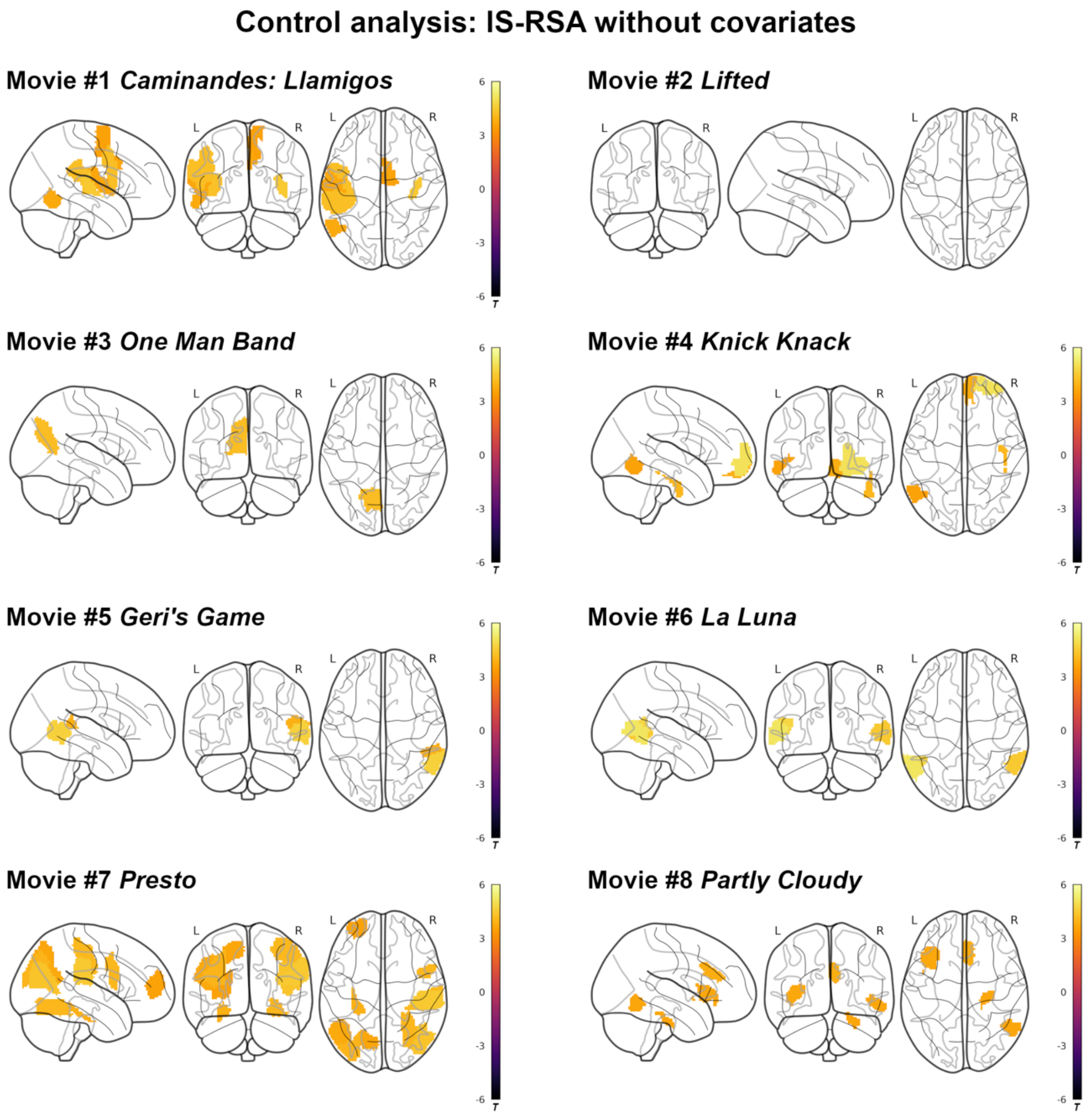


**Supplementary Figure 10. IS-RSA results without covariates in the CABB dataset.**

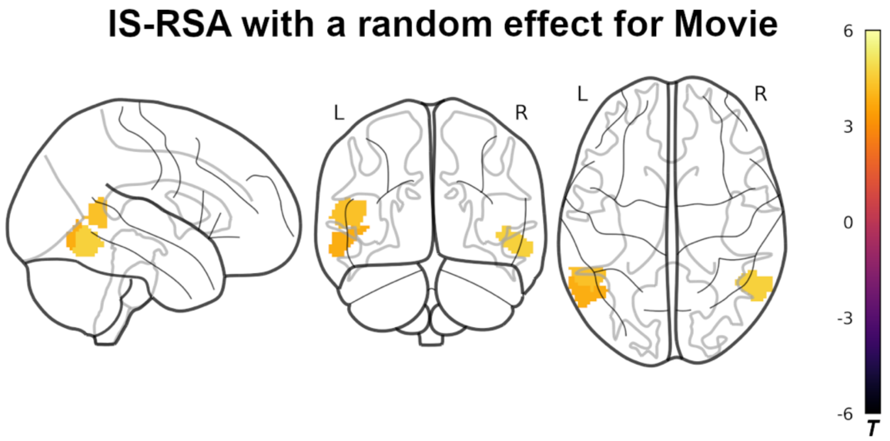


**Supplementary Figure 11. IS-RSA in the CABB dataset including a random effect for movie.** Brain regions where subject similarity in conceptual representations of novel objects was significantly associated with ISC, estimated using a linear mixed-effects model that included a random intercept for movie.

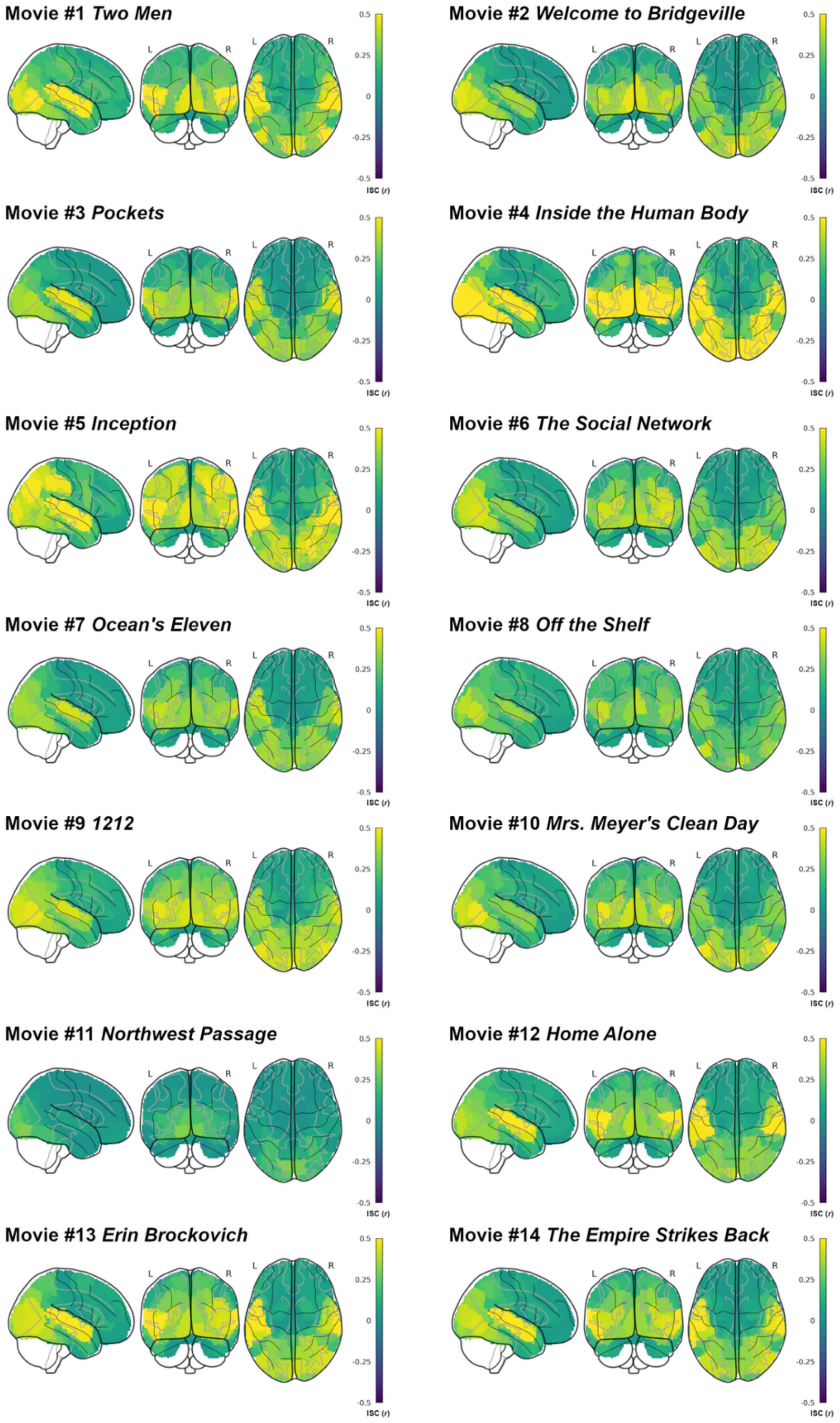


**Supplementary Figure 12. Inter-subject correlations for individual movies in the HCP dataset.** Spatial distribution and magnitude of inter-subject correlations (ISC) across the cortex, shown separately for each of the 14 movie clips.

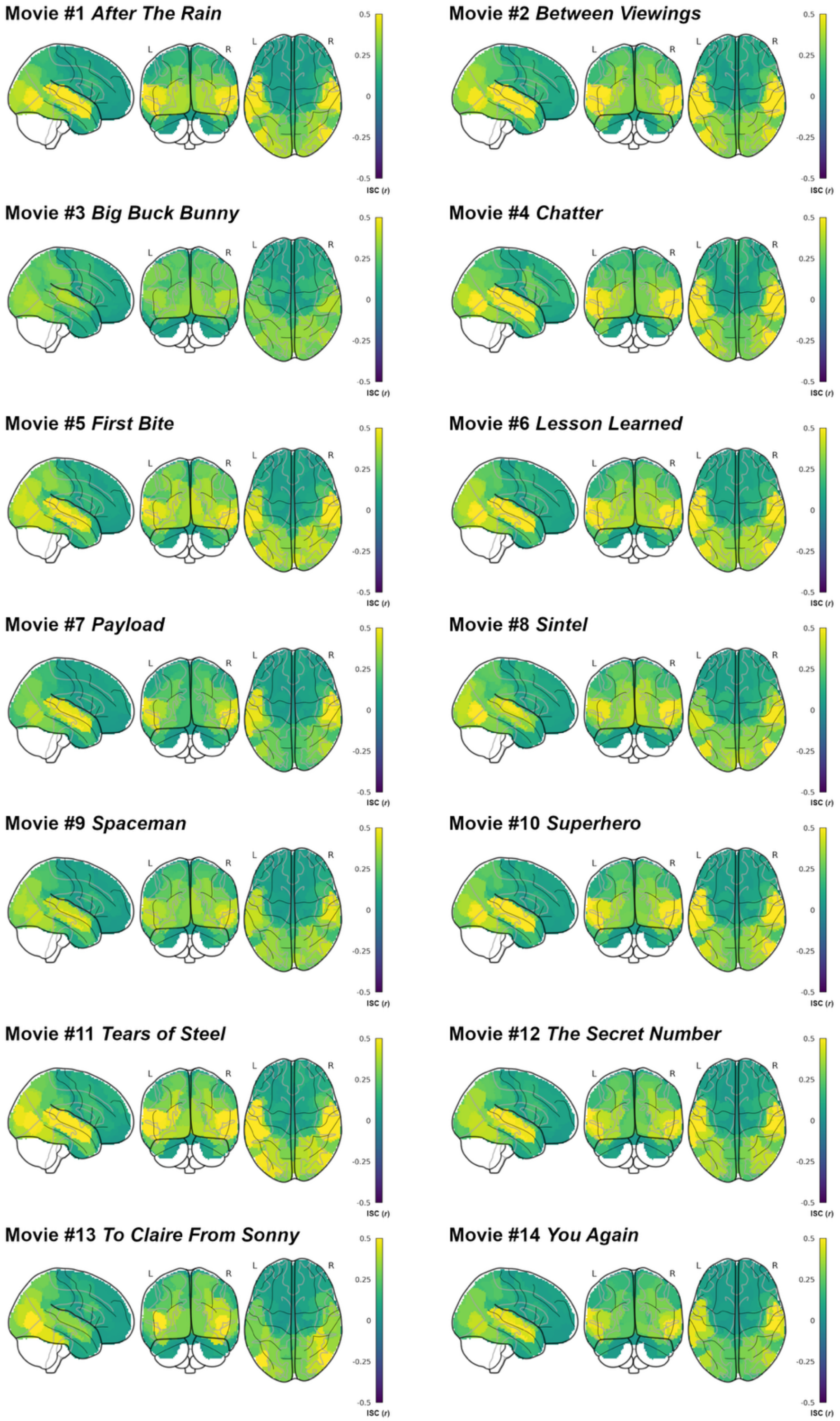


**Supplementary Figure 13. Inter-subject correlations for individual movies in the Emo-FilM dataset.** Spatial distribution and magnitude of inter-subject correlations (ISC) across the cortex, shown separately for each of the 14 movies.

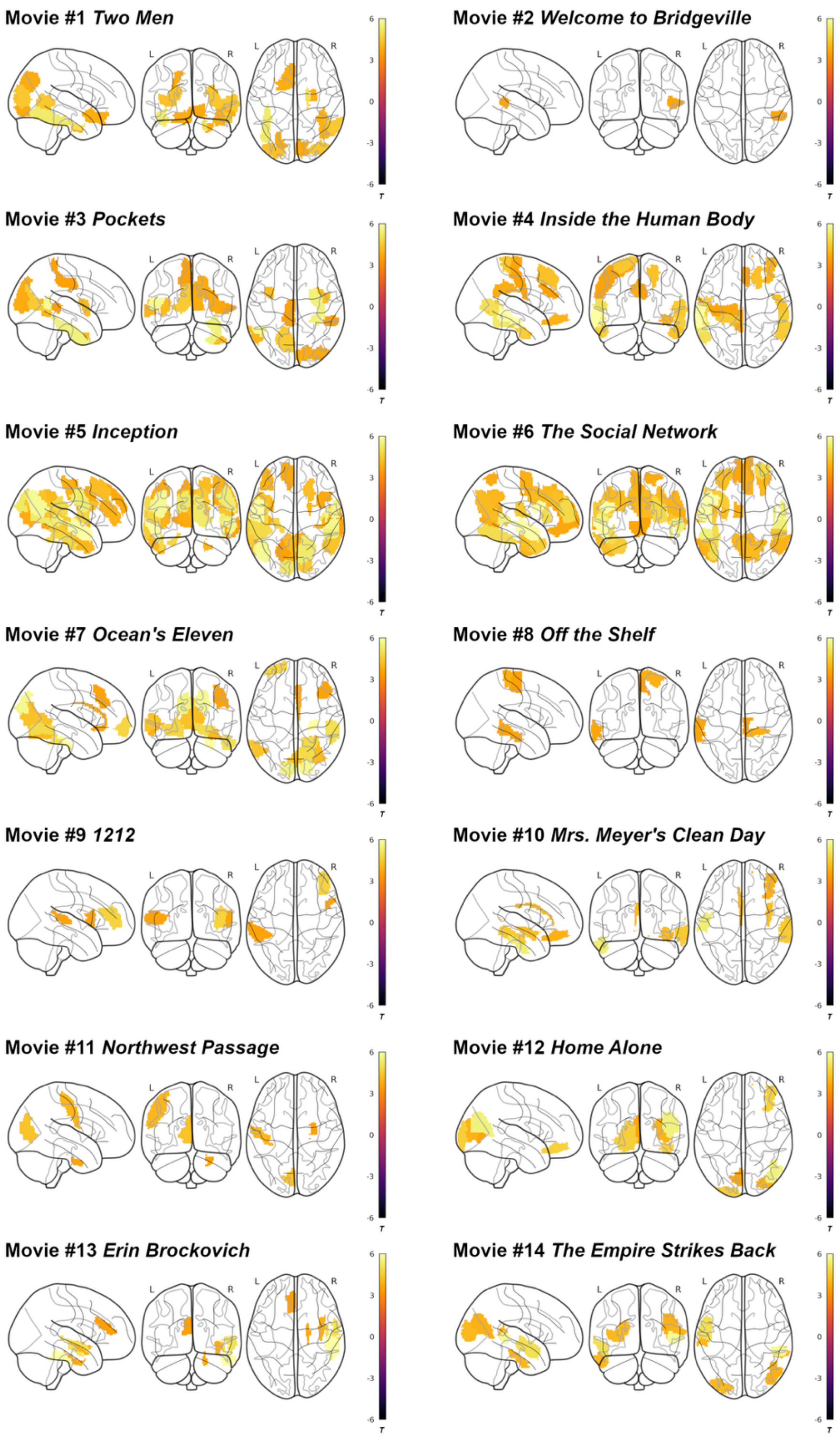


**Supplementary Figure 14. IS-RSA with Big Five personality traits in the HCP dataset.**

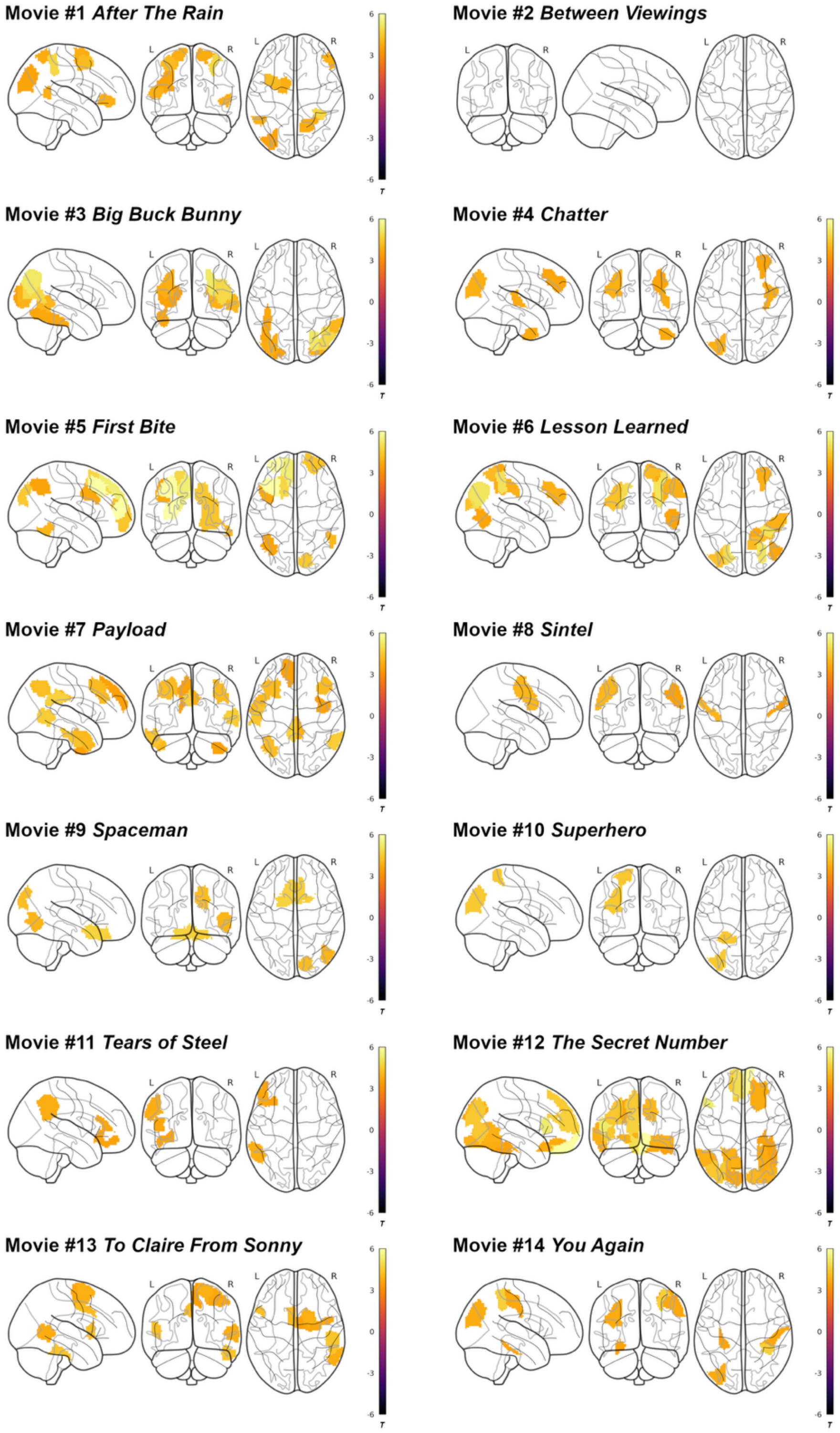


**Supplementary Figure 15. IS-RSA with Big Five personality traits in the Emo-FilM dataset**

### Supplementary Tables

**Supplementary Table 1. Set of features on which the novel objects were rated.**
In the Features task, subjects rated the novel objects on 29 features, provided here in the original Dutch with English translations. A lead-in sentence was displayed at the top of the screen (“In hoeverre zie je dit plaatje als …”; English translation: “To what extent do you view this image as …”). This table was adapted from Eijk et al. (2022).

| Feature (original Dutch) | Feature (English translation) |
| --- | --- |
| afgerond | rounded |
| puntig | pointy |
| symmetrisch | symmetrical |
| langwerpig | elongated |
| plat | flat |
| compact | compact |
| visueel complex | visually complex |
| aandacht vragend | demanding attention |
| iets met een kenmerkende kleur | something with a characteristic color |
| licht of helder om te zien | light or bright on the eyes |
| groot | large |
| klein | small |
| iets met een kenmerkende smaak | something with a characteristic taste |
| iets met een kenmerkende geur | something with a characteristic smell |
| makkelijk hoorbaar | easily hearable |
| gerelateerd aan beweging | related to movement |
| menselijk | human |
| iets met een hoofd/gezicht | something with a head/face |
| iets met een lichaam | something with a body |
| gerelateerd aan acties met het gezicht/de mond | related to actions with the face/the mouth |
| gerelateerd aan acties met de hand/arm | related to actions with the hand/arm |
| gerelateerd aan acties met de voet | related to actions with the foot |
| iets met een vaste plaats/locatie | something with a fixed place/location |
| iets waarvoor tijd (tijdstip of duur) relevant is | something for which time (time point or duration) is relevant |
| iets waar jij directe ervaring mee hebt | something you have direct experience with |
| iets wat jou of anderen helpt | something that helps you or others |
| iets waardoor je verrast wordt | something that you are surprised by |
| positief/plezierig | positive/pleasant |
| negatief/onplezierig | negative/unpleasant |

**Supplementary Table 2. Results of the IS-RSA for separate movies in the CABB dataset.** Associations between subject similarity in conceptual representations of novel objects, estimated in the Features task, and inter-subject correlation (ISC) during movie watching were quantified by linear mixed-effects models for each cortical parcel, while controlling for subject similarity in lexical representations, estimated in the Naming task, and sex and age. Parcels with statistically significant associations are ordered by parcel number, as defined by the Brainnetome atlas (Fan et al., 2016).

| Movie # | Parcel number | Brain region | $ß_{STD}$ | *t* | $p_{FWE}$ |
|---|---|---|---|---|---|
| 1 | 10 | SFG, Right Superior Frontal Gyrus A6m, medial area 6 | 0.05 | 3.70 | 0.02 |
| 1 | 63 | PrG, Left Precentral Gyrus A6cvl, caudal ventrolateral area 6 | 0.05 | 3.98 | 0.007 |
| 1 | 97 | ITG, Left Inferior Temporal Gyrus A37vl, ventrolateral area 37 | 0.05 | 3.67 | 0.03 |
| 1 | 145 | IPL, Left Inferior Parietal Lobule A40rv, rostroventral area 40 (PFop) | 0.05 | 4.02 | 0.006 |
| 1 | 163 | INS, Left Insular Gyrus G, hypergranular insula | 0.06 | 4.00 | 0.007 |
| 1 | 164 | INS, Right Insular Gyrus G, hypergranular insula | 0.06 | 4.10 | 0.004 |
| 1 | 184 | CG, Right [anterior] Cingulate Gyrus A24cd, caudodorsal area 24 | 0.05 | 3.60 | 0.03 |
| 3 | 151 | Pcun, Left Precuneus dmPOS, dorsomedial parietooccipital sulcus (PEr) | 0.05 | 3.52 | < 0.05 |
| 4 | 27 | MFG, Left Middle Frontal Gyrus A10l, lateral area10 | 0.05 | 3.62 | 0.03 |
| 4 | 28 | MFG, Right Middle Frontal Gyrus A10l, lateral area10 | 0.07 | 5.00 | < 0.001 |
| 4 | 48 | OrG, Right Orbital Gyrus A11m, medial area 11 | 0.06 | 3.76 | 0.02 |
| 5 | 86 | MTG, Right Middle Temporal Gyrus A37dl, dorsolateral area37 | 0.06 | 4.14 | 0.004 |
| 5 | 124 | pSTS, Right posterior Superior Temporal Sulcus cpSTS, caudoposterior superior temporal sulcus | 0.04 | 3.80 | 0.02 |
| 6 | 85 | MTG, Left Middle Temporal Gyrus A37dl, dorsolateral area37 | 0.07 | 5.31 | < 0.001 |
| 6 | 86 | MTG, Right Middle Temporal Gyrus A37dl, dorsolateral area37 | 0.06 | 4.57 | < 0.001 |
| 6 | 123 | pSTS, Left posterior Superior Temporal Sulcus cpSTS, caudoposterior superior temporal sulcus | 0.06 | 4.65 | < 0.001 |
| 7 | 19 | MFG, Left Middle Frontal Gyrus A46, area 46 | 0.06 | 3.99 | 0.007 |
| 7 | 64 | PrG, Right Precentral Gyrus A6cvl, caudal ventrolateral area 6 | 0.05 | 3.91 | 0.01 |
| 7 | 106 | FuG, Right Fusiform Gyrus A37mv, medioventral area37 | 0.05 | 3.98 | 0.007 |
| 7 | 113 | PhG, Left Parahippocampal Gyrus TL, area TL (lateral PPHC, posterior parahippocampal gyrus) | 0.05 | 3.55 | 0.04 |

| | | | | | |
|---|---|---|---|---|---|
| 7 | 127 | SPL, Left Superior Parietal Lobule A7c, caudal area 7 | 0.05 | 4.02 | 0.006 |
| 7 | 135 | IPL, Left Inferior Parietal Lobule A39c, caudal area 39 (PGp) | 0.05 | 4.12 | 0.004 |
| 7 | 136 | IPL, Right Inferior Parietal Lobule A39c, caudal area 39 (PGp) | 0.05 | 4.28 | 0.002 |
| 7 | 143 | IPL, Left Inferior Parietal Lobule A39rv, rostroventral area 39 (PGa) | 0.05 | 3.98 | 0.007 |
| 7 | 144 | IPL, Right Inferior Parietal Lobule A39rv, rostroventral area 39 (PGa) | 0.05 | 3.65 | 0.03 |
| 7 | 146 | IPL, Right Inferior Parietal Lobule A40rv, rostroventral area 40 (PFop) | 0.06 | 4.38 | 0.001 |
| 7 | 147 | Pcun, Left PrecuneusA7m, medial area 7 (PEp) | 0.04 | 3.68 | 0.02 |
| 7 | 160 | PoG, Right Postcentral Gyrus A2, area 2 | 0.05 | 3.97 | 0.008 |
| 7 | 199 | LOcC, Left lateral Occipital Cortex mOccG, middle occipital gyrus | 0.04 | 3.64 | 0.03 |
| 7 | 210 | LOcC, Right lateral Occipital Cortex lsOccG, lateral superior occipital gyrus | 0.04 | 3.56 | 0.04 |
| 8 | 37 | IFG, Left Inferior Frontal Gyrus A44op, opercular area 44 | 0.06 | 3.92 | 0.01 |
| 8 | 98 | ITG, Right Inferior Temporal Gyrus A37vl, ventrolateral area 37 | 0.04 | 3.59 | 0.04 |
| 8 | 112 | PhG, Right Parahippocampal Gyrus A35/36c, caudal area 35/36 | 0.05 | 3.96 | 0.008 |
| 8 | 180 | CG, Right [anterior] Cingulate Gyrus A32p, pregenual area 32 | 0.06 | 3.95 | 0.008 |

**Supplementary Table 3. Results of the IS-RSA across all movies in the CABB dataset.** Associations between subject similarity in conceptual representations of novel objects, estimated in the Features task, and inter-subject correlation (ISC) during movie watching were quantified by linear mixed-effects models for each cortical parcel, while controlling for subject similarity in lexical representations, estimated in the Naming task, and sex and age. A single model was fit across all movies, with movie included as a random effect. Parcels with statistically significant associations are ordered by parcel number, as defined by the Brainnetome atlas (Fan et al., 2016).

| Movie # | Parcel number | Brain region | $ß_{STD}$ | $t$ | $p_{FWE}$ |
|---|---|---|---|---|---|
| all | 97 | ITG, Left Inferior Temporal Gyrus A37vl, ventrolateral area 37 | 0.02 | 3.80 | 0.02 |
| all | 98 | ITG, Right Inferior Temporal Gyrus A37vl, ventrolateral area 37 | 0.02 | 4.68 | < 0.001 |
| all | 123 | pSTS, Left posterior Superior Temporal Sulcus cpSTS, caudoposterior superior temporal sulcus | 0.02 | 4.31 | 0.002 |
